# Microglial low-affinity FcγR mediates the phagocytic elimination of dopaminergic neurons in Parkinson’s disease degeneration

**DOI:** 10.1101/2024.07.04.602092

**Authors:** P.V. Casanova, E. Saavedra-López, I. Freitag-Berenguel, M. Usandizaga, P. Martínez-Remedios, C. Giménez-Montes, E. Clinton, M. Roig-Martínez, G.P. Cribaro, C Barcia

## Abstract

Microglia-mediated neuroinflammation contributes to dopaminergic (DA) neurodegeneration in Parkinson’s disease (PD). It is thought that microglial cells interact with DA neurons of the substantia nigra pars compacta (SNpc), the most vulnerable region in parkinsonian neuropathology, playing a critical role through the process of neuronal loss. However, the specific mechanisms by which microglial reactivity is triggered and exerts such a deleterious effect remains unclear. Experimental models of PD in mice have shown that phagocytosis plays an essential role in the elimination of degenerating neurons, suggesting that blocking this microglial function could be beneficial to preserve remaining DA neurons. In the present work, we pinpoint the role of the Fcγ receptor (FcγR) as a potential trigger of microglial phagocytosis in PD. We found that the FcγR is overexpressed in the degenerating SNpc in postmortem samples of PD patients, alongside histological indications of phagocytic microglia. Likewise, experimental models of PD also show increased FcγR expression, together with evidence of similar phagocytosis events. Most importantly, blocking FcγR *in vitro*, by using neutralizing antibodies, reduced the microglial-mediated elimination of DA cells. High resolution imaging revealed that the FcγR is involved in phagocytic synapse formation, and appeared polarized, and segregated to actin-rich protruding cups, when interacting one-on-one towards single target DA cells. Additionally, by inhibiting FcγR downstream actin-polymerizing signaling Cdc42, microglia were unable to eliminate DA cells. Finally, experiments *in vivo*, using an experimental model of PD in mouse, revealed that passive immunotherapy with FcγR neutralizing monoclonal antibodies, as well as inhibiting its downstream signaling Cdc42, protect DA neurons from elimination. These results indicate that the FcγR may be a critical factor inducing DA neuron phagocytosis in PD and suggest a novel immunotherapeutic strategy targeting microglia to preserve neuronal loss.

## Introduction

A substantial body of evidence points to microglial-mediated neuroinflammation as a contributing factor for PD dopaminergic (DA) neurodegeneration^1,2^. However, the mechanisms driving this response are still largely unknown^3^. The brain area undergoing the highest neuronal depletion, the substantia nigra *pars compacta* (SNpc), displays reactive microglia^4^, indicating the presence of persistent nerve degeneration. Disease-mediated reactivity of microglia, detected by the overexpression of human leukocyte antigen (HLA)-DR (nomenclature for human major histocompatibility complex II [MHC-II]), has been associated with the progression of DA degeneration in PD^4,5^. Most intriguingly, HLA-DR reactive microglia were also observed in the SNpc of accidentally induced parkinsonism by 1-methyl-4-phenyl-1,2,3,6-tetrahydropyridine (MPTP) in humans^6^, years after the neurotoxic insult, suggesting a perpetuation of the neuroinflammatory compound impacting DA degeneration. Parallel findings were also reported in MPTP experimentally induced parkinsonism in monkeys^7,8^ and mice^9^ further supporting the critical role of a microglial-mediated response in DA neurodegeneration. Use of the MPTP model has since been instrumental for a better understanding of the microglial response in PD^10^, although essential questions still remain unanswered regarding the precise participation of neuroinflammation in DA neuronal loss.

Microglial activation in parkinsonism appears to be interferon-γ (IFN-γ)-dependent^11^, which is associated with a pro-inflammatory profile. In fact, increased levels of IFN-γ have been detected in PD patients^12,13^, suggesting the establishment of a proinflammatory environment for microglia to be primed, and subsequently be overactivated, promoting the neurodegenerative process^14^. Our previous line of research, also performed in the MPTP mouse model of PD, has shown that microglial reactivity precedes DA neuronal elimination^11^. In this context, microglial cells physically interact with degenerating DA neurons, establishing phagocytic body-to-body contacts before neuronal loss^15^, which can be prevented by blocking microglial motility through the inhibition of Rho-associated protein kinase (ROCK), affecting the function of cell division control (Cdc)-42^15^.

However, the mechanisms that drive neuronal phagocytic elimination by microglia in the parkinsonian scenario remain to be determined. One of the plausible hypotheses is the occurrence of humoral immunity^16^. The presence of immunoglobulin G (IgG) binding to SNpc DA neurons of parkinsonian patients^16^, together with the presence of FcγR in neighboring microglia, suggest FcγR-IgG interaction as a potential trigger of their phagocytic capacity. Still, this premise has scarcely been explored. Previous studies, performed in mice lacking FcγR, and inoculated with PD-patient-derived IgGs, showed reduced microglial activation and DA neurodegeneration in comparison with their wild type (WT) counterparts, supporting a detrimental role of FcγR-IgG interactions in PD^17^. Other studies, also using FcγR-deficient mice, point to α-synuclein excess as the potential trigger of the humoral response inducing the pathological and deleterious activation of microglia^18^. Thus, these reports indicate that the FcγR appears to be a potential activator of microglial cells in this scenario. However, whether the FcγR plays a direct role in the phagocytosis of DA neurons in PD is yet to be determined.

In leukocytes, namely monocytes and macrophages, FcγR-mediated phagocytosis is regulated by Cdc42^19,20^. Thus, the macrophage-like characteristics of microglia indicate that they may follow parallel mechanisms. In this context, the binding to FcγR activates the downstream signaling Cdc42, directly involved in the actin cytoskeleton polymerization, and responsible for driving the cell motility and the formation of the phagocytic cup^21–23^. In the present work, we wanted to further explore the role of FcγRs in the parkinsonian pathology but focusing on the phagocytic capacity of microglia followed by Cdc42 signaling. First, we analyzed the topographic expression of low affinity FcγRs CD16 and CD32 in the SNpc of postmortem samples of PD patients, and then tested whether this feature was mimicked in the MPTP-induced model of PD in mice. Experiments in vitro, modeling the IFN-γ mediated pro-inflammatory scenario in microglial and DA cell co-cultures, revealed the involvement of low affinity FcγRs in the phagocytic interactions of microglia, and showed that either blocking the FcγR with neutralizing antibodies, or inhibiting its downstream signaling Cdc42, prevents DA cells elimination. Finally, passive immunotherapy using CD16/32 neutralizing monoclonal antibodies in MPTP parkinsonian mice protected DA neurons from elimination, suggesting a novel and viable potential therapeutic strategy for PD.

## Results

### Microglial phagocytic FCγRs are increased in the SNpc of PD patients

Because humoral immunity has been suggested as a potential factor in the pathogenesis of PD^16^, DA neurons bound to IgGs may facilitate the trigger of a FcγR-mediated response, inducing microglial phagocytosis^16^. To explore this, we analyzed the expression of FcγR in *postmortem* samples of PD patients. We first characterized the loss of neuromelanin-containing DA neurons of the SNpc of PD samples in comparison with non-parkinsonian individuals. Mesencephalic sections from PD patients showed clear depigmentation of the SNpc, especially visible in the ventrolateral region, as has been previously characterized histologically^24–27^ and most recently with single-cell genomic profiling^28^. Quantification of the neuromelanin-containing neurons revealed a significant degeneration of the SNpc in parkinsonian patients (**Supplementary figure 1 A, B**). A regional quantification, discriminating the dorsal and ventral tier of the SNpc showed a dramatic neuronal loss in the ventrolateral area (**Supplementary figure 1 A, C-E**). The characterization of the microglial-mediated neuroinflammation showed marked HLA-DR immunoreactivity in the SNpc of PD patients. Likewise, Iba-1 immunostaining revealed highly reactive microglia in the SNpc, both in nigrosomes and nigral matrix (**Figure 1A-C**). Close examination revealed Iba-1 microglial cells surrounding neuromelanin-containing neurons in PD patients (**Figure 1C**). Importantly, microglial cells adjacent to DA neurons showed evidence of phagocytosis. Higher magnification analysis in the SNpc of postmortem samples of parkinsonian patients revealed microglial cell branches displaying rounded terminal pouches, cellular structures compatible with ball-and-chain phagocytosis (**Figure 1D and E**). These events indicate the presence of moderate but chronic neuroinflammation.

**Figure 1.**
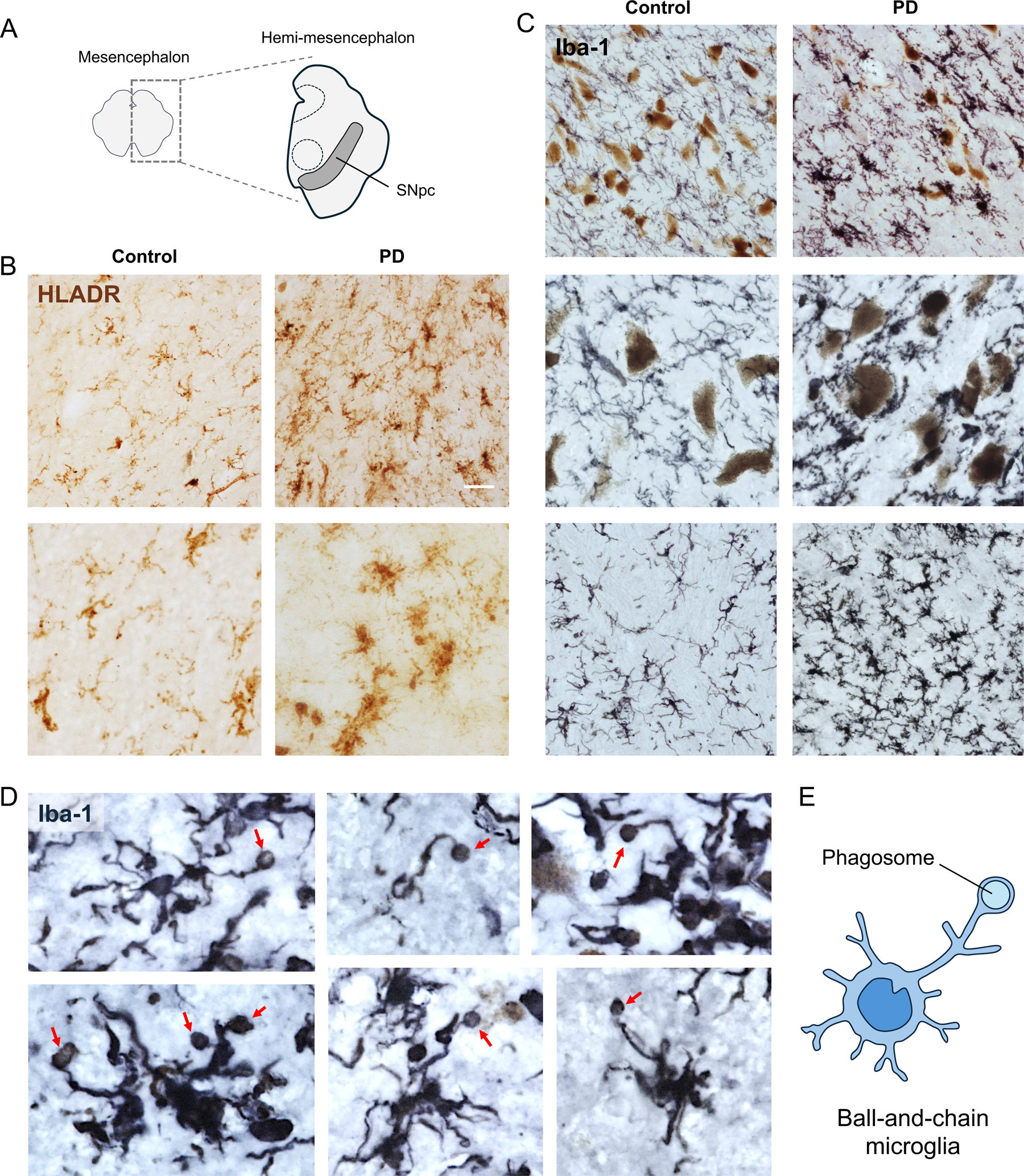
PD patients display hallmarks of microglial-mediated neuroinflammation and evidence of microglial phagocytosis. **A**. Diagram representing the anatomical area of study, section of hemi-mesencephalon, containing the SNpc. **B**. Immunoreactivity of HLA-DR was increased in the SNpc of PD patients compared with non-parkinsonian individuals (Top panel shows low magnification, and bottom panel displays higher magnification). **C.** Immunostaining for Iba-1 revealed reactive microglia in PD patients surrounding melanized DA neurons (Top panel with low magnification and middle panel with higher magnification). Note reactive microglia in close apposition to neurons. Bottom panel displays an image from nigral matrix to visualize the different morphology. **D.** Iba-1 labeling show microglia displaying rounded terminal buds (red arrows) compatible with ball-and-chain phagocytosis in the SNpc of PD patients. **E.** Schematic diagram of a ball-and-chain microglia.

Histological analysis of the expression of CD16 in the mesencephalon showed labelled microglial cells in the SNpc, with high immunoreactivity in PD patients (**Figure 2A**). Further examination revealed some CD16^+^ rounded cells, in perivascular areas and parenchyma resembling macrophages (**Figure 2B**). Histological assessment of CD32 showed very limited expression, only present in a few rounded cells within the SNpc, most likely compatible with perivascular macrophages or infiltrated monocytes (**Figure 2C**). The topographic examination of CD16^+^ microglia in the subregions of the SNpc showed higher reactivity in the dorsal region (**Figure 2D, E**). Importantly, stereological quantification of both degenerating areas of the SNpc revealed a significant increase of CD16 expressing microglia in PD patients but limited to the dorsal tier (**Figure 2E, F**). This pattern of expression is consistent with the spatiotemporal progression of the neuronal loss in PD, which begins in the ventral areas followed by the dorsal regions^25^. These results indicate that the humoral response, involving FcγR-expressing microglia, may be critical in the process of DA neurodegeneration, precisely in the places where the degeneration of remnant DA neurons may still be in place, potentially favoring their phagocytic elimination in PD.

**Figure 2.**
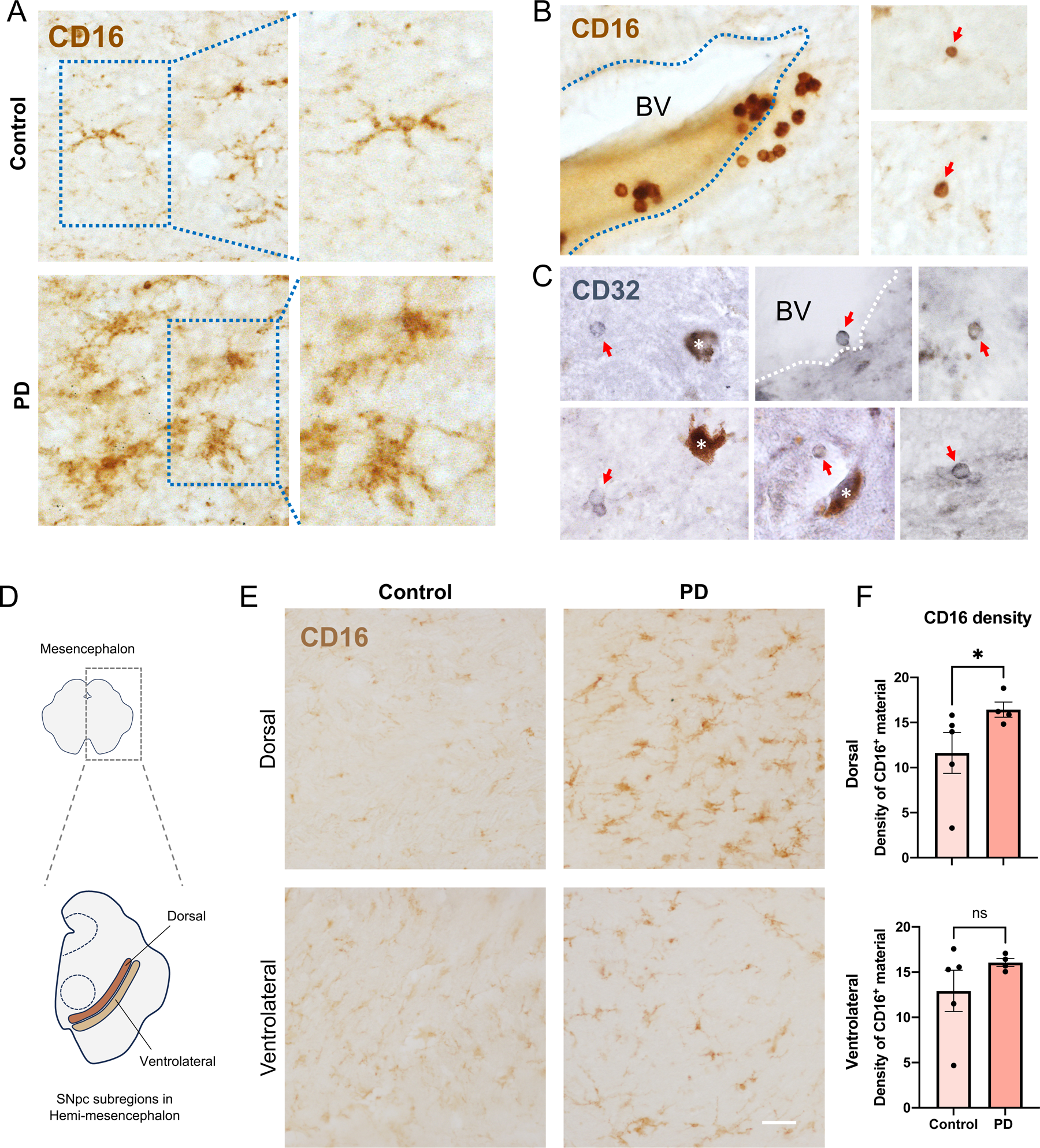
PD patients show increased CD16-expressing microglia in degenerating SNpc. **A.** Immunoreactivity of CD16 in the SNpc shows cells with microglial morphology in PD patients and controls. Some areas of the SNpc in PD patients display CD16^+^ microglia with reactive phenotype. **B**. CD16^+^ immunostaining also revealed rounded cells compatible with macrophages, surrounding blood vessels (BV) and in the parenchyma proper (red arrows). **C.** CD32 immunolabeling also shows cells with rounded morphology (red arrows), in BV, or in the parenchyma, sometimes in close apposition to neuromelanin-containing neurons (white asterisks). **D.** Diagram of a representative coronal section of a human mesencephalon at the level of the superior colliculus (top illustration), and the representative fixed hemi-mesencephalon used for the quantification, depicting the dorsal and ventrolateral tiers of the SNpc (bottom illustration). **E**. Representative images of the human tissue samples showing expression of CD16^+^ microglial cells in the dorsal and ventrolateral area of the SNpc in control individuals and PD patients. Note that microglial cells of the dorsal area of the SNpc from PD patients show higher CD16 immunoreactivity compared with control individuals. **F.** Quantification of CD16^+^ microglial cells density in the dorsal area of the SNpc shows a statistically significant increase in PD patients compared with controls (top graph). In contrast, the quantification of CD16^+^ density in the ventrolateral area did not show statistically significant differences between control individuals and patients. *p<0.05 compared to control. Scale bar in E: 100 μm.

### Microglial phagocytic receptor CD16/32 is increased in parkinsonian mice

As the SNpc of parkinsonian patients showed an increase of FcγR, to better understand the phenomenon in the parkinsonian scenario, we wanted to explore whether the expression of low affinity FcγRs could be replicated in a PD mouse model. We analyzed the SNpc of MPTP-injected mice by using antibodies against FcγR CD16/32 in sections of the mesencephalon. Concomitant with a 35% DA neuronal loss, immunolabeled sections showed CD16/32-expressing microglial cells in the mesencephalon of parkinsonian mice compared with controls (**Figure 3A**). Importantly, stereological quantification of CD16/32^+^ cells revealed a significant increase in the SNpc of MPTP-treated animals compared with saline injected animals, mimicking the increase seen in human disease. Although phagocytosis has previously been described by us as a critical event in the elimination of DA neurons in the MPTP model^15^, we wanted to corroborate the visualization of remaining phagocytic events in the SNpc. For this, we multi-labeled the sections to detect microglia, DA neurons of the SNpc, as well as the nuclei (**Figure 3C**). Detailed confocal analysis of these samples showed phagocytic events in the SNpc of MPTP-treated animals, characterized by the formation of large phagosomes close to the cell body (**Figure 3D-F**), as well as ball-and-chain structures (**Figure 3G-I**). In the latter, a phagocytic pouch formed at the tip of a microglial branch containing a pyknotic nucleus. Both formations suggest the phagocytic clearance of degenerating neurons after induced DA neurodegeneration, but the ball-and-chain phagocytosis suggests the continuation of the phagocytic process with moderate inflammatory response.

**Figure 3.**
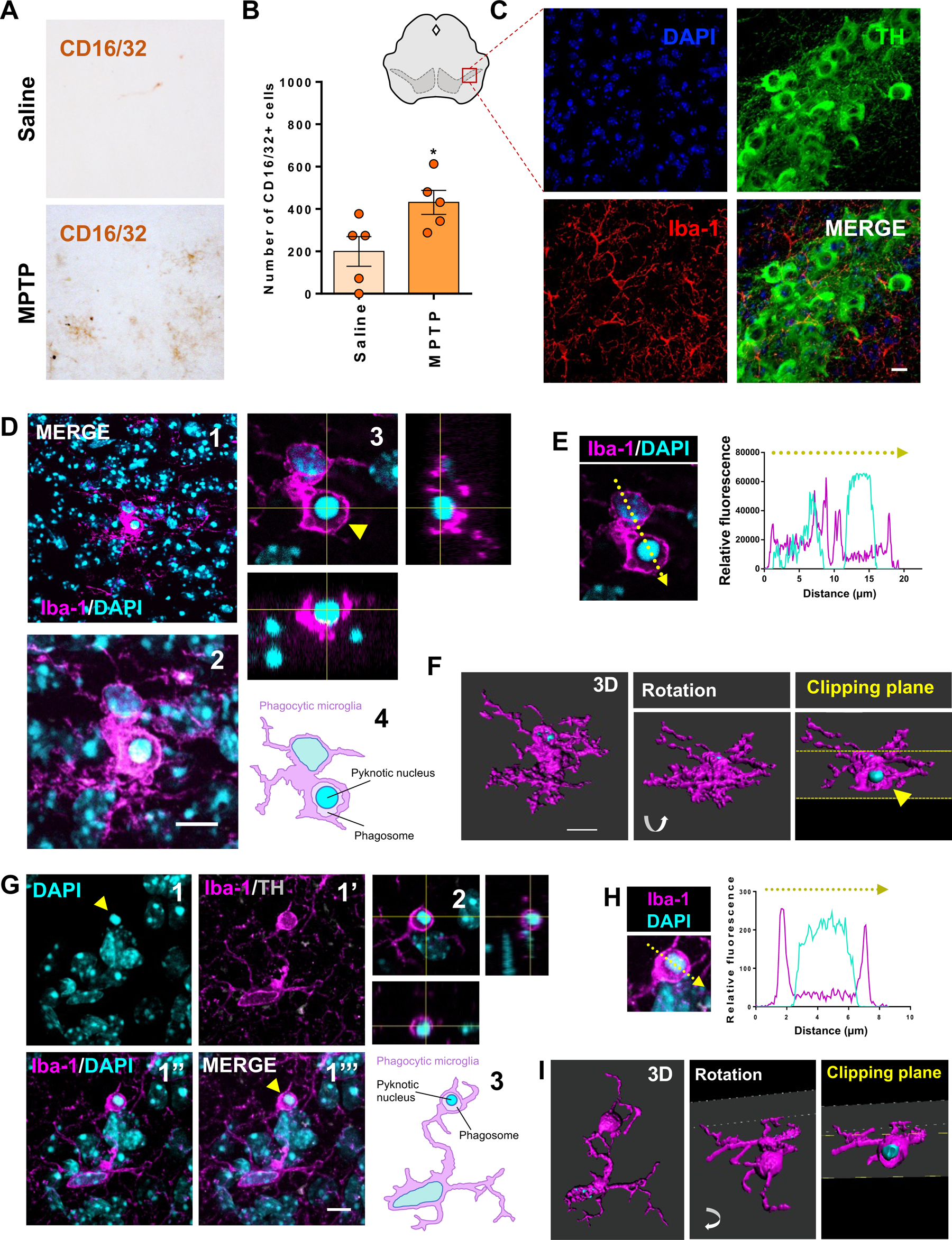
Microglial phagocytic receptor CD16/32 is increased in a mouse model of PD. **A**. MPTP-induced degeneration induces expression of CD16/32. Representative image of immunohistochemistry demonstrating the expression of CD16/32 in the MPTP-treated mice. **B**. Stereological quantification of the number of CD16/32 positive cells indicating a higher number of CD16/32 labeled microglia in the MPTP group compared with the control (* p˂0.05). **C.** Representative image obtained from mouse SNpc with microglia expressing Iba-1 (red), dopaminergic neurons expressing TH (green) and nuclei DAPI (blue) (Scale bar: 10 µm). **D**. Representative phagocytic event in the SNpc. A pyknotic nucleus, stained with DAPI inside Iba-1^+^ microglia can be appreciated in overview of maximum intensity projection of the z-stack (1) and with higher magnification (2). Single 0.5 mm optical slide and orthogonal views evidencing the phagosome (3) (yellow arrowhead). A schematic diagram of the phagocytosis event is shown (4) (Scale bar: 10 mm). **E**. Plot profile of relative fluorescence along the yellow arrow from image in D, showing higher fluorescence of DAPI in the pyknotic nucleus compared with the normal nucleus and the Iba-1 limits of the phagosome. **F**. 3D reconstruction from image in D showing the structure of the microglia and a clipping plane at the level of the phagosome showing the containing pyknotic nucleus (yellow arrowhead). **G.** High resolution stack image of a microglia cell with a long process containing pyknotic nucleus (yellow arrowhead). Microglia immunostained for Iba-1 (magenta), dopaminergic neurons immunostained for TH (grey), and nuclei counterstained with DAPI (cyan) (1,1’,1’’,1’’’). Single 0.5 mm optical slide and orthogonal views are shown evidencing the engulfing event along the z-axis and allowing the visualization of the pyknotic nucleus inside an Iba-1^-^ space, compatible with a phagosome (2) (Scale bar: 8 µm). A schematic diagram of the phagocytosis event is shown (3) **H**. Plot profile of relative fluorescence from yellow broken arrow delineated in the image, displaying the high fluorescence of DAPI consistent with the condensed pyknotic nucleus. High fluorescence of Iba-1 is also detected surrounding the pyknotic nucleus and limiting the phagosome. **I**. 3D reconstruction of ball-and-chain phagocytosis in the SNpc. Consecutive images show a rotated image with or without a clipping plane to show the inside of the phagosome containing the pyknotic nucleus.

### CD16/32 expression relates with canonical proinflammatory response *in vivo*

As the MPTP PD model induces CD16/32 microglial expression in the areas of neuronal degeneration, we wanted to test whether a sole proinflammatory stimulus was able to trigger CD16/32 expression and induce DA neuronal loss. For this, we used LPS as the inducer of the proinflammatory reaction in the brain. In this case, because direct injection of LPS in the SNpc causes severe parenchymal damage^29^, we used an indirect model, injecting LPS in the striatum but visualizing the effect on the SNpc (**Figure 4A**). Histological analysis showed an increase of microglial reactivity, evidenced by Iba-1 expression, both in striatum and SNpc, restricted to the injected hemisphere, in contrast with the contralateral side (**Figure 4B-C**). In addition, CD16/32 immunoreactivity was also expressed in microglia of the injected hemisphere (**Figure 4D**). Interestingly, quantification of DA neuronal loss and microglial reactivity revealed a significant neuronal loss in the ipsilateral hemisphere, in contrast with a transient downregulation of TH in the contralateral side (**Figure 3E-F**). This effect on neuronal loss was concomitant with a peak of microglial reactivity, especially evident three days after LPS injection in the ipsilateral side but being mitigated seven days post injection (**Figure 4E-F**). Most interestingly, detailed confocal analysis of the SNpc also showed evidence of ball-and-chain phagocytic events, also suggesting the phagocytic clearance of degenerating DA neurons during the resolution of a pure proinflammatory scenario (**Figure 4G**). These results show that LPS-and MPTP-induced parkinsonism share similar proinflammatory responses, involving CD16/32 expression and phagocytic clearance.

**Figure 4.**
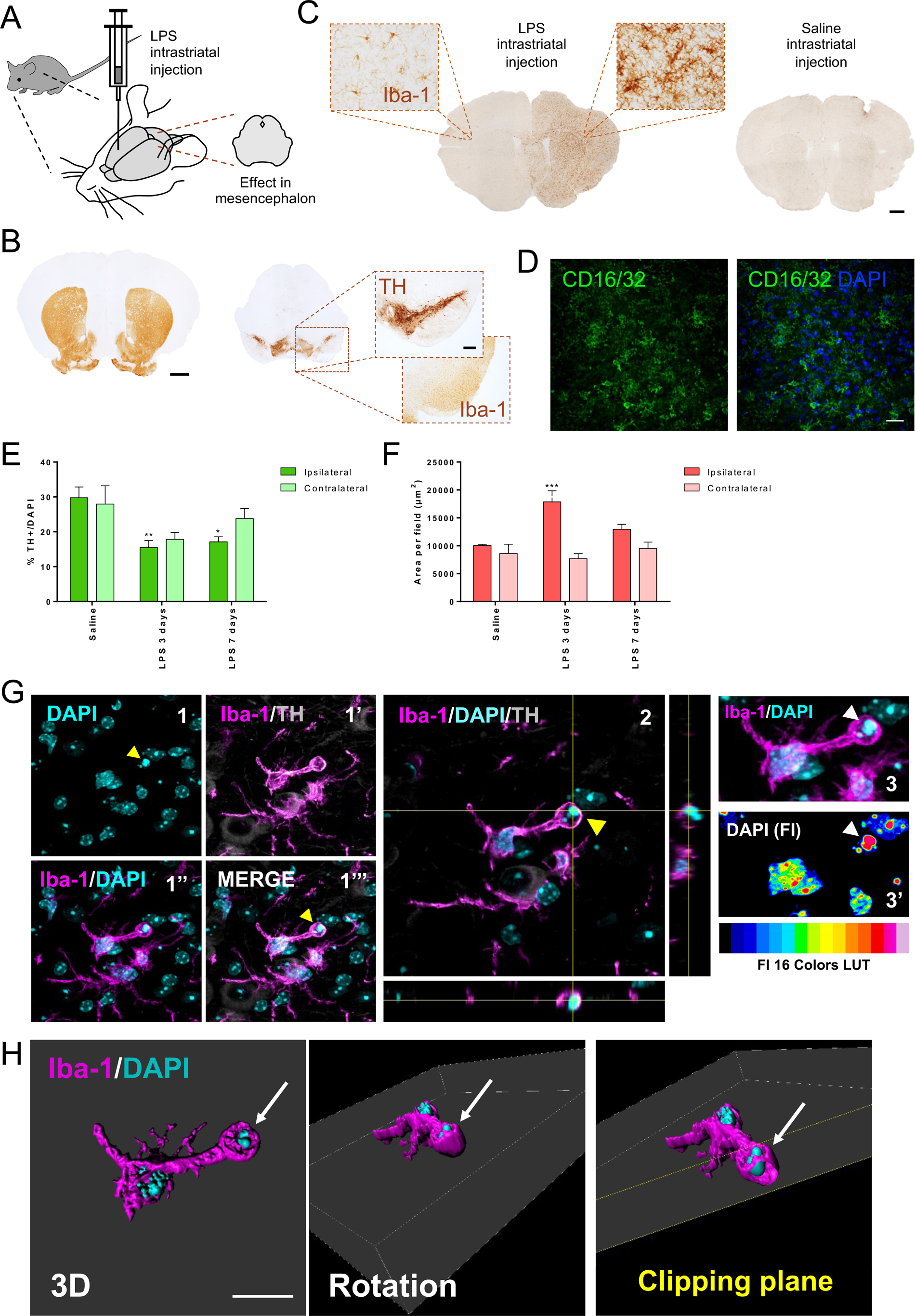
Proinflammatory insult in the nigrostriatal pathway induces phagocytic CD16/32 expressing microglia. **A** Diagram of the experimental procedure. **B**. Representative image of the striatum with dopaminergic fibers stained for TH (Scale bar: 500 µm). Mesencephalon with dopaminergic neurons of the SNpc stained for TH and Iba-1 (scale bar: 200 µm). **C.** Representative image of mice striatum with high expression of Iba-1 in microglia (and potentially macrophages at the site of injection) in the ipsilateral hemisphere. Right panel displays a control striatum injected with saline showing low expression of Iba-1 (scale bar: 500 µm). **D.** Confocal image representing expression of CD16/32 in the nigrostriatal pathway after intrastriatal injection of LPS (scale bar: 20 µm). **E.** Quantification of the number of dopaminergic neurons of the SNpc expressing TH. Significant decrease of neurons is appreciated at three days and seven days after LPS injection in the ipsilateral hemisphere (** p˂0.01 and * p˂0.05 vs. saline). **F.** Quantification of the area of Iba-1 expressed in microglia in the SNpc indicated significant increase at three days post-injection in the ipsilateral hemisphere. (*** p˂0.001 vs. contra and ipsilateral saline, vs. LPS three days contralateral and LPS seven days contralateral). **G.** Phagocytic event with “ball-and-chain” structure in the SNpc. High-resolution stack image of a microglial cell with a branch containing a pyknotic nucleus (yellow arrowhead). Microglia immunostained for Iba-1 (Magenta), dopaminergic neurons for TH (grey) and nuclei counterstained with DAPI (cyan). Single 0.5 mm optical plane and orthogonal views at the central plane evidenced the engulfing event along the z stack, showing the pyknotic nuclei, with highly condensed DNA inside an Iba-1^−^ space, consistent with a phagosome structure. Fluorescence intensity analysis with a 16-color LUT, showed high intensity levels in pyknotic fragmented nucleus inside the phagosome (3 and 3’’). **H**. 3D reconstruction of phagocytic microglia from panel G in zenithal view and rotated in perspective. Clipping plane (white arrow) showing the phagosome containing the pyknotic nucleus (Scale bar: 10 µm).

### CD16/32 expression relates with canonical proinflammatory response *in vitro* and specific 1:1 interactions

To better understand the factors and mechanism underlying the phagocytic properties of microglia in the context of DA degeneration, as well as to disentangle CD16/32 function in this scenario, we set up an in vitro model combining microglia and DA cells in 2D co-cultures. Utilizing the BV2 microglial cell line, we first characterized this model in terms of the level of activation and phagocytic capacity. We used LPS and IFN-γ as the canonical stimulus to achieve a proinflammatory response. The level of activation was first measured by nitrite release and Iba-1 expression. We observed a very significant increase in nitrites when microglia were treated with the combination of LPS and IFN-γ (both 24 and 48h after the treatment) (**Supplementary figure 2 A, B**). A slight increase was seen when IFN-γ was used alone (**Supplementary figure 2 A, B**), and interestingly, induced a relevant surge of cellular proliferation (**Supplementary figure 2 C-G**). As indicated, in addition to nitrites, the level of activation was also measured by the expression of Iba-1, being higher in accordance with increasing concentrations of either IFN-γ or IFN-γ/LPS (**Supplementary figure 2 H-J**). LPS and IFN-γ concentrations of 100 ng/mL 0.5 ng/mL, respectively, were determined as the optimal combination, reaching the highest level of activation but without exerting major cellular toxicity, to use for subsequent experiments.

To analyze the potential phagocytic capacity of activated microglia after the canonical activation with LPS/IFN-γ, we quantified the number of PC12 dopaminergic cells and the microglia-target interactions. We observed a significant reduction after 24h when microglia were activated (**Supplementary figure 3 A-E**). However, the number of interactions was not significantly modified between 4 and 24 hours. Importantly, detailed confocal analysis revealed microglial cells containing TH reactive material or pyknotic nuclei, indicating the phagocytic capacity of microglia (**Supplementary figure 3 F-J**). Quantification of these events revealed a significant increase in phagocytosis when microglial cells were activated, indicating the induction of the phagocytic capacity of microglial cells after canonical proinflammatory stimulus. Knowing that higher activation of microglial cells was achieved when stimulating with the combination of IFN-γ and LPS than with IFN-γ or LPS alone, we analyzed the effect of the sequential stimulation in different orders. In this case, stimuli were separated into primary or secondary, changing the order of the administration in every condition, and waiting 24 hours between one and the other. Finally, 24 hours after the last stimulus, activation was analyzed. To study which of the stimuli generates the maximum response, and the highest activation as a consequence, we used both BV2 microglia. Our results indicated that reactivity of microglial cells takes place with both options. But most importantly, the maximal activation was reached when the microglial cells were first stimulated with IFN-γ and later challenged with LPS (**Supplementary figure 4**). Because sequential stimulation with IFN-γ and LPS induced the maximum microglial activation, we set up subsequent co-culture experiments with BV2 and PC12 cells with this procedure to mimic the neuroinflammatory environment in PD where microglia and DA neurons interact. BV2 were first stimulated with IFN-γ and 24 hours later challenged with LPS, then, 24 hours later PC12 cells were added to the culture. Immunofluorescence labeling allowed the quantification of cell-to-cell contacts and the visualization of phagocytic events.

As we wanted to analyze the process of microglial phagocytosis, and keeping in mind that cell-target contacts may take place prior to the analyzed time window (four hours after activation), we decided to explore shorter timepoints to study intercellular interactions. In this set of experiments, following IFN-γ priming, canonically activated microglia significantly increased the expression activation markers Cd11b and F4/80, but most importantly, induced a highly significant increase in CD16/32, suggesting the production of FcγR to prepare for the phagocytic process (**Figure 5A, B**). Microglia-DA cell contacts were analyzed 10, 20, 30 and 60 minutes after co-culture initiation, showing an increasing number of interactions from 10 up to 30 minutes. Then, a drop in interactions occurred at 60 minutes, concomitant with the significant elimination of DA cells (**Figure 5C-D**), indicating the importance of the intercellular contacts in this process. In fact, in this *in vitro* set up, besides a high number of intercellular appositions, evidence of phagocytic elimination was also seen (**Figure 5E**). To further analyze the specificity of the microglia-target interactions, we characterized different types of contacts in 2D co-cultures. After corroborating the elimination of DA cells 1h after interaction, only when microglia were canonically activated (**Figure 5F, G**), we assessed the dynamics of intercellular contacts, according to the number of DA cells being contacted (**Figure 5H**). In each case, the number of contacts increased in the first 30 minutes, being 1:1 contacts being the most prominent interactions (**Figure 5I**). However, the number of interactions appeared to be similar regardless of the activation state. The analysis in terms of percentage revealed very similar levels of microglia-target interactions despite whether DA cells were presented to naïve or activated microglia (**Figure 5J**). Analysis of the 30-minute peak showed that 1:1 interactions were significantly prominent in all three treatments (**Figure 5K**). Conversely, the analysis of the 60-minute drop showed fewer 1:1 contacts in canonically activated microglia, concomitant with the phagocytic elimination of dopaminergic cells (**Figure 5K**). Importantly, these results suggest that microglia participate in the elimination of cellular targets on a 1:1 basis, in a specific and polarized manner-dealing with one target at a time.

**Figure 5.**
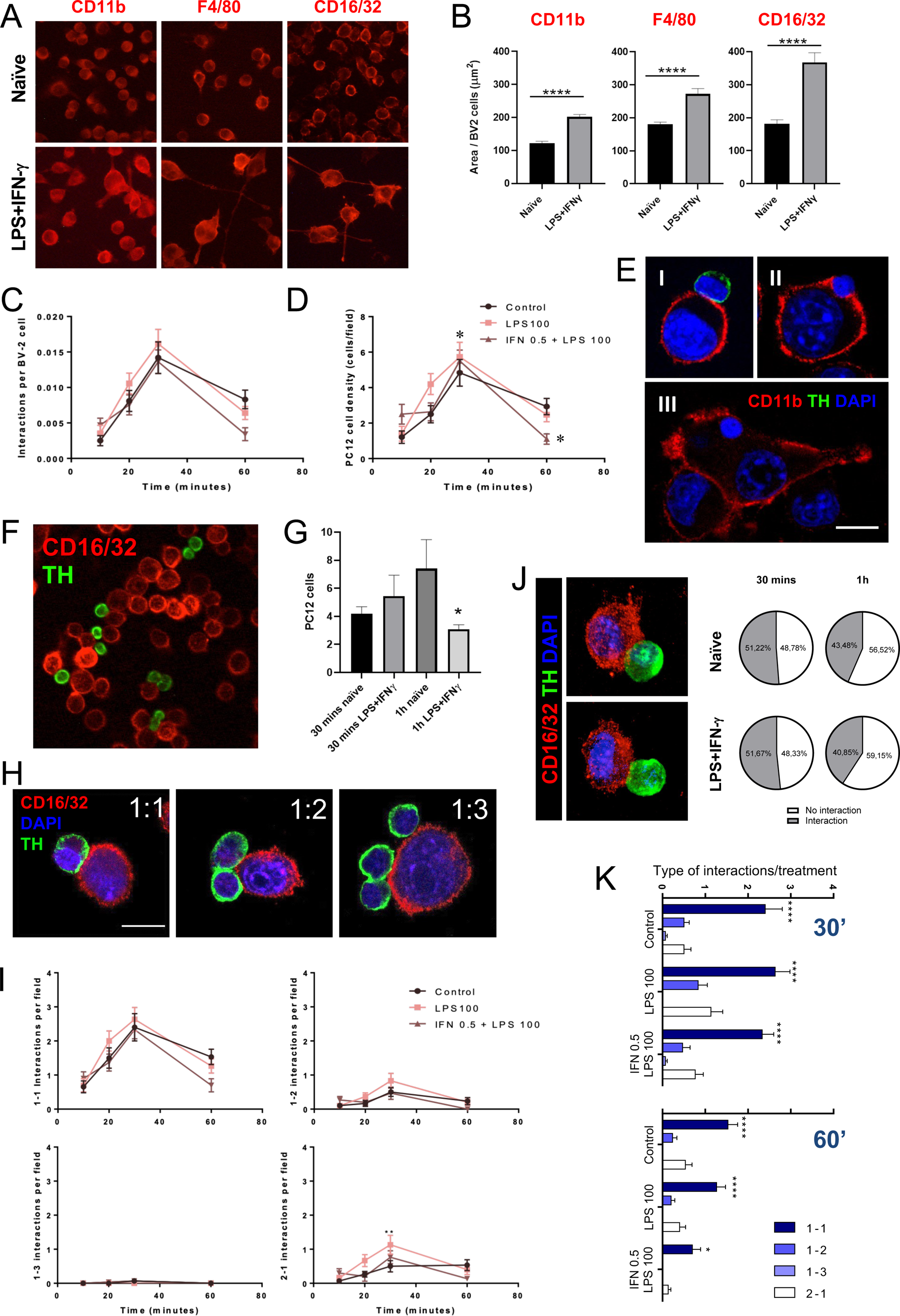
Microglial cells prominently interact with single dopaminergic cell targets. **A.** Representative microscopy images of non-activated (naïve) and activated microglia (LPS+IFN-γ). We used antibodies against CD11b, F4/80 and CD16/32 markers to detect the expression and morphology, and consequently, the level of activation of microglial cells after LPS/IFN-γ treatment. **B.** The quantification of the immunoreactivity expressing area of the three markers in BV2 microglial cells show a consistent increase after activation, especially high for phagocytic receptor CD16/32 (**** p<0.0001). **C.** The number of interactions between PC12 and BV2 cells varied with the same pattern through time in all the conditions with higher slope in activated microglia. **D.** PC12 cell density decreased significantly at 60 minutes when microglial cells were canonically primed (* p˂0.05 IFN-γ 0.5 + LPS vs. LPS100 and Control). **E.** Confocal images illustrating a characteristic contact (I) between BV2 microglial cell stained for Cd11b (red) and PC12 cell stained for TH (green). (II) BV2 (red) forming a phagocytic cup engulfing a pyknotic nucleus stained with DAPI (blue). (III) BV2 cell (red) containing a pyknotic nucleus (blue) inside the cell. Scale bar: 10 µm. **F.** Representative micrograph of the co-culture combining TH^+^ dopaminergic cells (green) and CD16/32^+^ microglia (red). **G.** Elimination of PC12 dopaminergic cells after one hour of interaction with LPS/IFN-γ-activated microglia. The quantification of PC12 cells in different conditions showed a significant decrease after one hour of interaction between naïve and LPS/IFN-γ-treated microglia (* p<0.05). **H.** Representative images of types of interaction between microglia and dopaminergic neurons in co-cultures. **I.** Quantification of interactions, according to different types (1-1, 1-2. 1-3 and 2-1), within the first hour after co-culture initiation. **J.** Representative images of 1-1 interactions, and pie charts showing the percentage of microglial cells establishing intercellular interactions with dopaminergic cells. **K.** Statistical analysis of types of interactions 30 and 60 minutes after the initiation of the co-culture. Note that 1-1 contacts are prominent at both time points. (* p<0.05, **** p˂0.0001 vs. every other treatment).

### CD16/32 segregate at the phagocytic synapse and mediates dopaminergic cells elimination

Considering this prominent 1:1 polarization, we then analyzed the microanatomy of intercellular contacts between microglia and DA cells, focusing on CD16/32 receptors. We observed that the expression of CD16/32 was lower at the center of the interface, but higher at the periphery, suggesting a segregation of the receptor during the formation of the phagocytic synapse (**Figure 6A-C**). By using high resolution confocal microscopy, single synapsing interactions were imaged in 3D and the relative fluorescence was analyzed at the intercellular interface. The analysis showed that the expression of CD16/32 fluorescence at the center of the phagocytic synapse was significantly lower and accumulated at the peripheral rim, suggesting CD16/32 participation in the establishment of the phagocytic synapse towards the elimination of DA cells (**Figure 6D**). To demonstrate the functionality of this phagocytic process, we blocked CD16/32 with neutralizing antibodies in microglia-dopaminergic cell co-cultures. We treated the cells with CD16/32 neutralizing antibodies either before or after the induction of the pro-inflammatory stimulus (**Figure 6E**). In both cases, a significant reduction of DA elimination was achieved (**Figure 6F**), demonstrating that CD16/32 are fundamental receptors in the process of elimination of target cells by reactive microglia.

**Figure 6.**
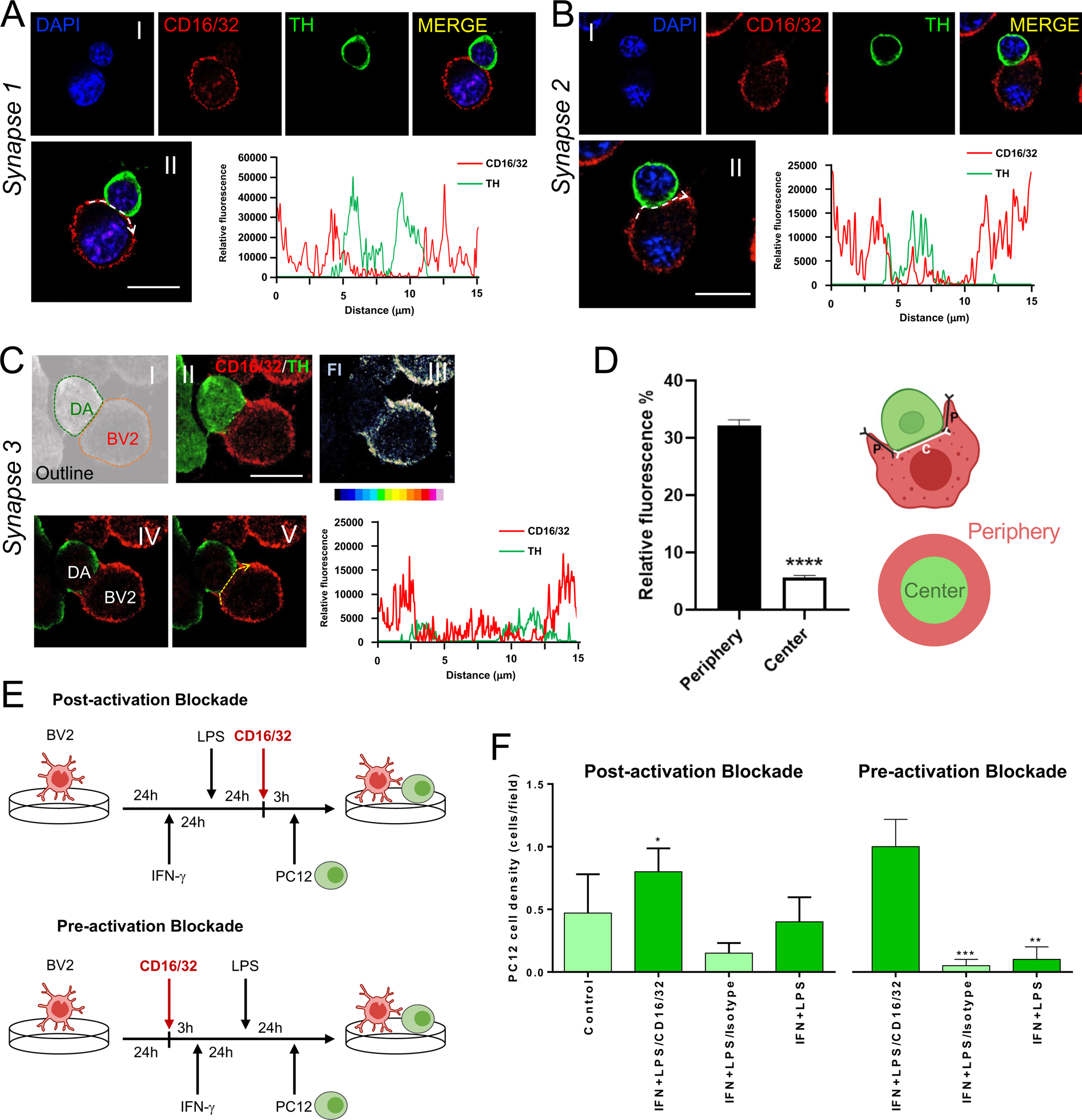
Blocking CD16/32 with neutralizing antibodies prevents microglial-mediated phagocytic elimination of dopaminergic cells *in vitro*. **A.** (*Synapse 1*, top panel [I]) Representative confocal image of a CD16/32^+^ BV2 microglia interacting with a TH expressing PC12. (Synapse 1, bottom panel [II]) Plot profile of relative fluorescence measured at the synapsing interface from image II (white broken arrow) displaying high fluorescence levels of the phagocytic receptor CD16/32 at the border of the interface, starting point of the phagocytic-cup, and low levels coinciding with the contact of the PC12 dopaminergic cell. **B.** (*Synapse 2*, top panel [I]) Representative example of interacting microglia and dopaminergic cell as displayed in A. (*Synapse 2*, top panel [II]) Plot profile analysis as performed in A. **C.** (Synapse 3, top panel) Grayscale of a dopaminergic cell (DA) and a microglial cell (BV2) outlined with colored broken lines (I). Maximum projection of CD16/32 and TH markers showing the arranging a phagocytic cup by BV2 microglia (II). Image for 16-color based relative fluorescence of the CD16/32 marker from image II shows accumulations at the border and segregation at the center (III). Scale on the bottom; white represents the highest value and black the lowest. (Synapse 3, bottom panel) Middle optical plane of the phagocytic synapse (IV). Plot profile of relative fluorescence, obtained from image V (yellow broken arrow) showing high expression of CD16/32 in the phagocytic cup surrounding the PC12 cell but low expression in the center where the contact with the BV2 cell is established. Scale bars equal 10 μm in all phagocytic synapse images. **D.** Quantification of the relative fluorescence shows high CD16/32 levels at the periphery of phagocytic cup compared with remarkable low levels of expression coinciding with the contact with the target cell (**** p<0.0001). Diagrams of the cell-cell interaction (top) and interface (bottom) are shown. Segments “P” (periphery) and “C” (center) indicates the measured interface resulting in a CD16/32-rich peripheral ring as opposed to CD16/32-low center. **E.** Diagram of the experimental procedure in vitro to block CD16/32 with neutralizing antibodies, either before or after the inflammatory stimulus. **F.** Quantification of DA cells after the treatment with neutralizing antibodies. In both cases, when BV2 microglia were blocked with CD16/32 neutralizing antibodies, PC12 dopaminergic cells were protected from phagocytic elimination. *p<0.05 (IFN-γ + LPS / CD16/32 significant against every other treatment) **p<0.01, *** p˂0.001 (against IFN-γ + LPS / CD16/32).

### Inhibiting CD16/32 downstream signaling Cdc42 in vitro prevents dopaminergic cells from elimination

Because FcγR-mediated phagocytosis is dependent on Cdc42 signaling^19,20^, we wanted to test whether its pharmacological inhibition prevents the elimination of DA cells. Moreover, Cdc42 is directly related with F-actin cytoskeletal polymerization and the formation of the phagocytic cup, driving the phagocytosis process. In this context in vitro, we first wanted to analyze the cellular distribution of FcγR expression in microglial cells. Labeled microglia showed essentially membranous labeling of CD16/32. Importantly, kinetic microglial cells showed accumulations of CD16/32 in the leading lamella and trailing uropod, a characteristic distribution, similar to F-actin fibers, when cells are undergoing motile trajectories (**Figure 7 A-C**). On the other hand, Cdc42 expression was also visualized in microglia, displaying mostly cytoplasmic accumulations but especially high when close to F-actin rich areas, as protrusions and lamellas (**Figure 7D, E**). Phagocytic events, containing mature phagocytic cups engulfing DA cells showed dense accumulations of F-actin at the periphery of the interface accompanied by a segregation at the center (**Figure 7F**). This distribution was analogous to the arrangement of CD16/32, also observed in intercellular microglia-DA cells contacts, verifying the mechanistic interaction of both components and their cooperative participation in the phagocytic process. Pharmacological inhibition of Cdc42 was implemented by using ML141 diluted in DMSO. First, to rule out any major toxicity of the inhibitor or vehicle, we performed a viability test comparing with H_2_O_2_ by using different ML141 concentrations. We did not record any major toxicity when using either the inhibitor or the vehicle at the chosen doses (**Figure 7G**). Importantly, inhibiting Cdc42 prevented the elimination of DA cells when presented to canonically activated microglia (**Figure 7H**) both when Cdc42 inhibition was done before or after microglial activation (**Figure 7H**).

**Figure 7.**
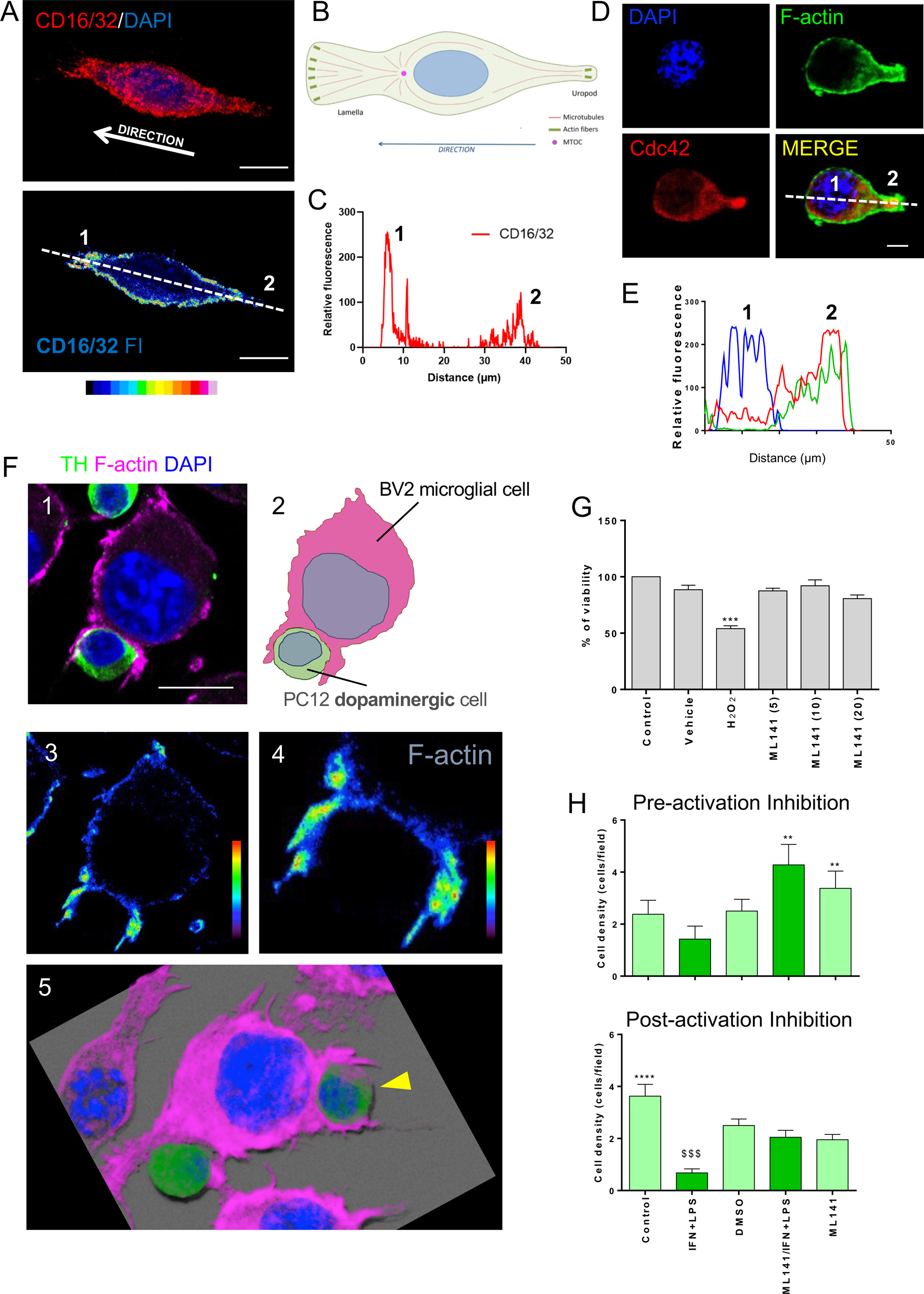
Inhibiting CD16/32 downstream signaling Cdc42 prevents microglia-mediated DA elimination. **A.** CD16/32 accumulates at the leading front of motile microglia. Top panel shows maximum intensity projection of a BV2 cell marked with CD16/32 and DAPI. Leading lamellas (left pole) and the uropod (opposite pole) indicate its direction. Bottom panel shows a 0.5 mm optical plane represented in a scale of 16 colors. Scale on the bottom (white represents the highest value and black the lowest). Leading front lamellas (1) and uropod (2) are indicated. **B.** Illustration of characteristic motile microglial cell showing essential cytoskeletal elements (Modified from Roig-Martinez et al., 2019)^43^. **C.** Plot of relative fluorescence of motile microglia measured (white broken line) from image on the left. Note the higher fluorescence at the leading front (1) and a smaller peak at the uropod (2). All images scale bars equal 10 μm. **D.** Representative image of a BV2 microglial cell expressing high density of Cdc42 (red) in a F-actin-rich protrusion. The nucleus was counterstained with DAPI (blue). Scale bar: 3 μm. **E.** Plot profile of relative fluorescence displays high expression of Cdc42 corresponding with high fluorescence of F-actin at the protruding edge (2) compared with the nucleus (1). **F.** Confocal image of microglial cell establishing a phagocytic cup around a dopaminergic cell (1). The PC12 DA cell is marked with dopaminergic neuronal marker, TH (green), and the microglial cell is stained by F-actin (magenta). (2) Diagram of the engulfing process depicted in 1. (3) Image of F-actin fluorescence intensity of the engulfing microglia seen in 1. Note the higher intensity is displayed at the borders of the phagocytic cup. (4) higher magnification of the phagocytic cup to visualize the characteristic accumulations of F-actin. Scale bar: 10 mm. (5) 3D reconstruction of the phagocytic event displaying the microglia arranging the F-actin rich phagocytic cup around a dopaminergic cell (yellow arrowhead). **G.** Increasing concentrations of ML141 were safe for BV2 cell viability. MTT cell viability assay for BV2 microglial cells submitted to increasing concentrations of ML141 was performed to test their safety: 500 μg of H_2_O_2_ resulted in approximately 50% decrease of the cell viability. But no significant decrease was appreciated after the administration of ML141 or vehicle (*** p˂0.001 H_2_O_2_ significant against every other treatment). **H.** Quantification of the number of PC12 DA cells at 60 min. of interaction with BV2 microglia. BV2 were treated either before or after activation with Cdc42 inhibitor ML141. The proinflammatory-mediated elimination of DA cells was prevented when BV2 cells were incubated with ML141. (Pre-activation inhibition ** p˂0.01 IFN-γ + LPS vs. ML141/IFN-γ +LPS and ML141. Post-activation inhibition $$$ p˂0.001 IFN-γ + LPS vs. every condition and **** p˂0.0001 control vs. IFN-γ + LPS).

### Passive immunotherapy with monoclonal antibodies blocking CD16/32 in the MPTP-induced PD model in mice prevents the elimination of DA neurons

As we showed that the elimination of DA cells was prevented *in vitro* by blocking CD16/32 and its downstream signaling Cdc42, we wanted to test this in a model of PD in mouse *in vivo*. For this, we used the acute model of MPTP injection, able to induce a detectable neuronal loss in the SNpc with a transient activation of microglial cells^11,15^. We blocked CD16/32 before the MPTP injection, following the most effective timing procedure seen above *in vitro*, and we quantified the DA neuronal loss and the microglial response in the SNpc (**Figure 8A**). Most importantly, passive immunotherapy by the treatment of monoclonal CD16/32 neutralizing antibodies was able to prevent the MPTP-induced elimination of DA neurons in the SNpc (**Figure 8A-C**). Conversely, 3D reconstructions of microglial cells displayed a reactive phenotype, in MPTP-treated animals, characterized by the increase in cell body size, and expanded branching (**Figure 8D**). Quantification of Iba-1 expressing area, showed a slight increase in parkinsonian animals, being less prominent in CD16/32 treated group (**Figure 8E**). Mirroring the previous experiment performed in vitro, we also analyzed the effect of inhibiting the CD16/32 downstream signaling Cdc42. First, we tested the effectiveness of the pharmacological inhibitor ML141 in reducing LPS-induced inflammatory response in the brain, specifically cortex and striatum. The treatment with Cdc42 inhibitor ML141 was able to suppress the LPS-mediated microglial inflammatory response in both areas (**Supplementary figure 5**), thus being an effective agent for taming microglial reactivity in this scenario. Using the same strategy, we analyzed the DA neuronal loss and microglial response in the MPTP model of PD, after the inhibition of Cdc42 (**Figure 8F**). We observed that the treatment with ML141 prevented the MPTP-induced elimination of DA neurons (**Figure 8G, H**), and concomitantly, reduced the microglial reactivity (**Figure 8G and I**).

**Figure 8.**
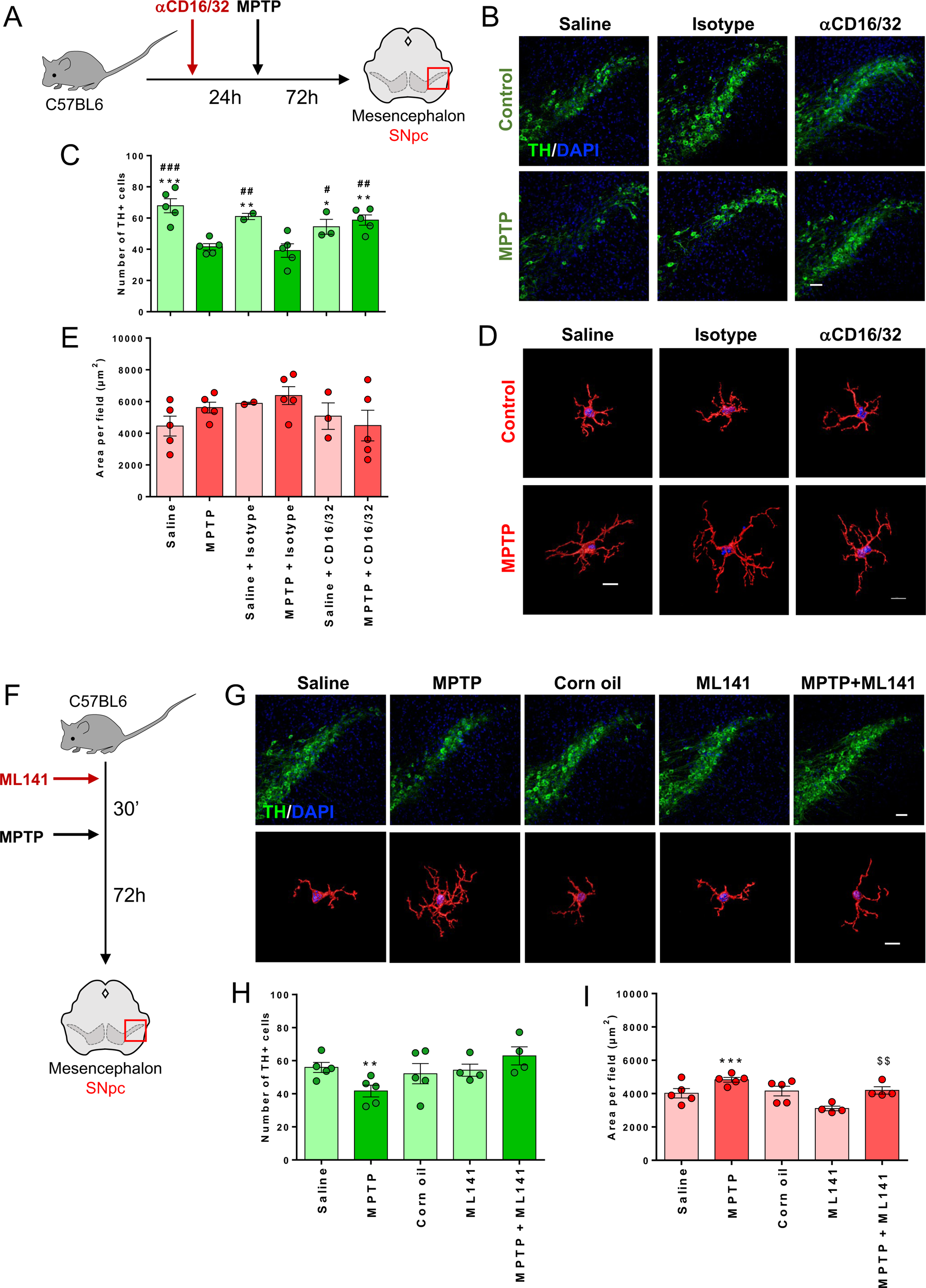
Inhibition of CD16/32 and its downstream signaling Cdc42 prevents dopaminergic elimination in MPTP-induced PD model. **A.** Diagram of the procedure used for the CD16/32 passive immunotherapy. Neuropathological histology was performed in the SNpc (red square). **B.** Representative confocal images showing the SNpc labeled with TH and counterstained with DAPI. Depletion of DA neurons can be appreciated in MPTP-treated animals. However, a protective effect can be seen in mice treated with CD16/32 neutralizing monoclonal antibodies (aCD16/32) (Scale bar: 30 mm). **C.** Quantification of the DA neurons in the SNpc demonstrating a significant decrease in the MPTP group which is prevented by the administration of CD16/32 (*** p˂0.001 MPTP vs. every treatment except MPTP + Isotype, ### p˂0.001 MPTP + Isotype vs. every treatment except MPTP). **D.** 3D reconstruction of representative microglial cells expressing Iba-1 in different treatments of the experiment (Scale bar: 10 µm). **C.** Quantification of the area of Iba-1 in the SNpc, indicating variation of microglial size when the animals were intoxicated with MPTP. **F.** Diagram of the procedure used for Cdc42 inhibition. **G.** Representative confocal images of histological analysis. Upper panel: Representative images of TH positive neurons of the SNpc in every treatment (scale bar: 30 µm). Lower panel: 3D reconstructions illustrating morphology and changes of size in microglial cells expressing Iba-1 (scale bar: 10 µm). **H.** Quantification of TH positive neurons indicating a significant decrease in the MPTP group and prevention by the previous administration of Cdc42 inhibitor ML141 (** p <0.01 with respect to saline). **I.** Quantification of the Iba-1 expressing area, which represents changes in microglial size. Microglial activation is present in the MPTP group evidenced by the increased area of Iba-1, which is decreased in the animals intoxicated with MPTP but previously treated with ML141 (*** p<0.001 with respect to controls, $$ p<0.01 with respect to ML141).

## Discussion

Our results indicate that the activation of microglial FcγR may underlie the phagocytic elimination of DA neurons in the PD neuroinflammatory context. The increase of FcγR in the degenerating areas of PD patients suggests that this response is ongoing during the pathological process. Thus, it is interesting to notice that the ventral tier of the SNpc, the area with the most dramatic DA neuron loss, does not show this increase in comparison with the dorsal tier, in which a higher population of DA neurons remains. This result indicates that FcγR response may be dependent on the presence of remnant neurons, still being active in the dorsal area, but no longer active in the ventral area after a major DA depletion. Consequently, this pattern of CD16-expression is coherent with the spatiotemporal progression of DA neuronal loss in PD, in which the most vulnerable population of neurons, located in the ventral SNpc^27^, is the first to be lost, followed by the dorsal regions in more advanced cases^25,26^. These results support the hypothesis that humoral immunity may be involved in PD pathogenesis, most likely by promoting phagocytosis through the interaction of microglial cells with IgG-bound DA neurons^16^, following the spreading of the pathology. In fact, this is consistent with the presence of reactive microglia surrounding remaining DA neurons and the evidence of ball-and-chain phagocytosis events.

This increase in FcγR reactive microglia is mimicked in parkinsonian mice after the neurotoxic insult by MPTP, and as a part of the neuroinflammatory response, further supporting the potential involvement of the phagocytic process. As previously reported, phagocytosis of DA neurons in the acute MPTP model peaks 48 h after the neurotoxin insult, with the maximum number of body-to-body, cell-cell interactions around this time-window^15^. However, when DA neuronal loss becomes evident, occurring after 72h, the number of phagocytic events is significantly reduced. In the present work, our 3D confocal analysis revealed remaining phagocytic events at that time point, 72 h after the insult, when the neuroinflammatory response is decreasing. In this context, we were able to image a ball-and-chain phagocytic type, consistent with the elimination of apoptotic corpses at the end of the acute MPTP-induced neurodegeneration and neuroinflammatory response. These structures have been seen previously in other scenarios, such as the clearance of apoptotic newborn cells in neurogenesis in the developing cortex^30,31^ or amygdala^32^ by unchallenged microglia. In the context of the parkinsonian scenario, this indicates that the phagocytic process may still take place after the nearly resolved inflammation. Identifying ball-and-chain phagocytosis in the MPTP mouse model parallels the phagocytic process previously described in MPTP-chronic parkinsonian monkeys^33^, and, most importantly, the phagocytic events seen here in PD patients.

Our in vitro experiments shed light on the mechanism of microglial phagocytosis in the context of parkinsonian neuroinflammation. Microglia appear to function on a one-on-one basis, each microglial cell physically interacting with a single DA cell target, and most interestingly, rearranging FcγR at the phagocytic synapse. This rearrangement requires the polarization of the microglial cell towards a single target. The intercellular engagement allows the communication between the cells, triggering the segregation of FcγRs to the peripheral area of the cell-cell interface. This reorganization of FcγR towards the protruding extensions of the intercellular interaction follows the pattern of the F-actin accumulations, suggesting the direct implication in the phagocytic process. This rearrangement seen here in microglia appears to be analogous to the pattern described for macrophages and neutrophils during FcγR-mediated phagocytosis^34^, where receptors and actin fibers are concentrated in the advancing cup-shape extensions. Importantly, neutralizing FcγR protects DA cells from microglial-mediated elimination, indicating its implication in the phagocytic process, potentially hampering the formation of the phagocytic cup. Most importantly, analogous results are seen in the mouse experimental model of PD, where DA neurons are preserved from elimination in animals treated with passive immunotherapy with monoclonal antibodies against FcγR.

The stimulation of FcγR activates the downstream signaling Cdc42 involved in the process of actin polymerization, driving the motility of the cell and phagocytosis. Particularly, Cdc42 is directly involved in the formation of the filopodia and lamellipodia^35–37^, key protruding elements established during cell polarization as well as the formation of the phagocytic cup^34^. In fact, these are essential actin-dependent cellular structures for microglial surveilance^38^, likely to participate in the process of phagocytosis in vivo. Therefore, impeding the actin polymerization, and the formation of filopodia and lamellipodia with Cdc42 inhibitor ML141 results in similar protection of DA cells in vitro. Analogously, the use of Cdc42 inhibitor in vivo, also protects DA neurons from elimination, suggesting that obstructing the microglial-mediated phagocytic process may be beneficial to preserve from parkinsonian degeneration.

Our results suggest that the use of immunotherapy targeting FcγR, tuning down the phagocytic capacity of microglia, could be beneficial to preserve DA neurons from elimination in PD. In the last decade, several protocols of immunotherapy, using monoclonal antibodies against amyloid-β (Aβ), have been developed and tested in clinical trials for Alzheimer’s disease. Because of the capacity of monoclonal antibodies to reach the brain parenchyma through transcytosis and via the glymphatic system, these therapeutics are delivered either intravenously, subcutaneously, or intraperitoneally, to bind and clear the Aβ pathological accumulations^39,40^. Similarly, immunotherapies for PD have also been developed with monoclonal antibodies against pathological accumulations of α-synuclein^41^ instead. Although the outcome of these therapeutic strategies is still conflicting, the emerging technologies show promise, not only to target misfolded proteins but also other cellular targets. In the present study we also propose the use of immunotherapeutic tools but, in this case oriented towards microglia as a cellular target to reduce the neuroinflammatory component involved in the pathology. Targeting the FcγR and its downstream signaling, would potentially prevent not only phagocytic elimination of DA neurons but also microglial motility, affecting their deleterious reactivity in the PD pathology.

## Methods

### Tissue samples from Parkinson’s disease patients

Samples and data from patients included in this study were provided by the Biobank *Banco de Tejidos CIEN* (PT17/0015/0014), integrated in the Spanish National Biobanks Network and they were processed following standard operating procedures with the appropriate approval of the Ethics and Scientific Committees. Particularly, this project was approved by the Fundación CIEN Scientific Committee (Reference code, S21005) and by the Ethics Committee on Human and Animal Experimentation (CEEAH) of the Universitat Autònoma de Barcelona (Reference code, CEEAH 5605). For conducting this study, postmortem mesencephalic brain areas, containing the SNpc, were obtained from patients diagnosed with PD, along with matching control subjects without parkinsonian diagnosis. Experimental grouping, age, sex, and clinical assessment were provided by the Biobank (**Table 1**). Brain tissue blocks of the hemi-mesencephalon were dissected in a coronal orientation, except for one case (**Table 1**), and fixed with formaldehyde at the Biobank. Samples were cryoprotected on arrival with cryoprotective solution (30% sucrose in 0.1M PB, pH: 7.4) and stored at -20°C until sectioning. Twelve series of 40 μm-thick sections of each tissue block were cut in the cryostat (CM3050S, Leica, Wetzlar, Germany) following the coronal orientation. Serial sections were then preserved at -20°C immersed in antifreeze solution (Ethylene glycol, glycerol in 0.1M PB, pH: 7.4) until processed for neuropathology staining and immunolabeling.

**Table 1.** Human samples.

| **Biobank Code** | **Sex** | **Age** | **Group** | **Diagnosis** | **Brain area** | **Section orientation** | **Fixative** |
| --- | --- | --- | --- | --- | --- | --- | --- |
| BCPA835 | M | 79 | PD | PD | Mesencephalon | Coronal | Formaldehyde |
| BCPA981 | M | 89 | PD | PD | Mesencephalon | Longitudinal/Transverse | Formaldehyde |
| BCPA1030 | M | 82 | PD | PD | Mesencephalon | Coronal | Formaldehyde |
| BCPA893 | F | 90 | PD | PD | Mesencephalon | Coronal | Formaldehyde |
| BCPA871 | F | 78 | PD | PD | Mesencephalon | Coronal | Formaldehyde |
| BCPA870 | M | 71 | CTRL | Non-PD | Mesencephalon | Coronal | Formaldehyde |
| BCPA732 | M | 57 | CTRL | Non-PD | Mesencephalon | Coronal | Formaldehyde |
| BCPA972 | M | 86 | CTRL | Non-PD | Mesencephalon | Coronal | Formaldehyde |
| BCPA701 | M | 55 | CTRL | Non-PD | Mesencephalon | Coronal | Formaldehyde |
| BCPA718 | F | 48 | CTRL | Non-PD | Mesencephalon | Coronal | Formaldehyde |

### Animals and surgery procedures

All the experimental procedures performed on animals were approved by the institutional CEEAH (Reference codes, CEEAH 1977 and CEEAH 2375) and by the local government (Reference codes, 10130 and 10123).

#### Intraperitoneal administration of LPS

For this experiment 25 male mice from the strain C57BL/6 were purchased from a commercial provider (Janvier Labs) at an age of three weeks. The animals were kept and maintained at the UAB animal house facilities in standard conditions, distributed in five groups of five mice per cage, and every cage was labeled with an experimental condition: saline, corn oil (Sigma Aldrich; Saint Louis, MO, USA), LPS, ML141, or ML141/LPS. First, the solutions were prepared fresh to follow the administration the same day. ML141 was diluted in corn oil up to a concentration for a 10 mg/Kg dose, and LPS was diluted in saline solution to a concentration for a 500 µg/Kg dose. The same volumetric amount of corn oil, in which ML141 was diluted (104 µL), or the same volumetric amount of saline, in which LPS was diluted (100 µL), were injected per animal in control groups, Corn oil and Saline respectively. On day 1, Corn oil group animals were administered (with a syringe with hypodermic needle) with 104 µl of corn oil, Saline group animals received 100 µl of saline solution, animals of the ML141 group received a 10 mg/Kg dose of ML141 diluted in corn oil, and the ML141/LPS animals were only administered 10 mg/Kg dose of ML141. On day 2, animals of the LPS group were injected with saline and 30 minutes later with a 500 µg/Kg LPS dose. The animals of the Corn oil group were injected with a dose of corn oil, and 30 minutes later with another dose of corn oil. Animals of ML141 group received the second dose of ML141 10 mg/Kg, the Saline group received a first dose of saline and a second one 30 minutes later. In the case of ML141/LPS, animals received a dose of 10 mg/kg of ML141 and 30 minutes later 500 µg/Kg of LPS. After 24 hours, animals were euthanized according to the guidelines of the CEEAH. Mice were anesthetized with a mixture of ketamine (Imalgene, Merial Laboratorios, Barcelona, Spain), at a dose of 100 mg/kg; and Medetomidine (Medetor, Ecuphar, Barcelona, Spain), at a dose of 0.5 mg/kg, then were perfused intracardially with 50 mL of Tyrode’s solution to achieve a complete blood perfusion and clearance. This was followed by a second perfusion phase with 50 mL of 4% PFA to obtain tissues fixation. The brain was then extracted and post-fixed in 4% PFA during 48 h. After post fixation, brains were stored in PBS and NaN_3_, until sectioning.

#### Intracranial injection of LPS

For this experiment, we also used three-week-old C57BL/6 male mice (n = 10) purchased from the same provider (Janvier Labs). Animals were kept and maintained in the UAB housing facilities with standard conditions. Each mouse was labeled on the tail for each experimental condition: eight animals for LPS and two animals for saline. Animals were first anesthetized with a mixture of anesthesia and analgesia: ketamine at a dose of 100 mg/kg; and medetomidine, at a dose of 0.5 mg/kg. LPS at a concentration of 1 mg/mL was inoculated stereotaxically in the striatum at specific coordinates: anteroposterior (AP): + 0.5 mm; mediolateral (ML): + 2.1 mm; dorsoventral (DV): - 3 mm, according to Bregma. After surgery, mice were injected with atipamezol (Antisedan, Ecuphar, Barcelona, Spain) at a dose of 2.5 mg/kg for medetomidine reversal. Once mice recovered, they were returned to the cage in normal conditions with normal feeding and maintained until the euthanasia. The same procedure was done with a control group, injecting intra-cranially 0.5 µL of saline solution. The animals were euthanized three or seven days after the intracranial injection (n = 5 and 3, respectively) by perfusion following the protocol described for the intraperitoneal administration of LPS in the previous section. Brains were extracted from the perfused mice and post-fixed in tubes with 4% PFA during 48 h. After post fixation, brains were stored in PBS and NaN_3_, until sectioning.

#### MPTP intoxication and Cdc42 inhibition

For the induction of the experimental model of PD, male mice (n = 25) with C57BL/6, a MPTP susceptible background, were acquired at three weeks of age (Janvier Labs). The animals were maintained in special disposable plastic cages in a room isolated from the UAB Animal House, with adequate conditions for MPTP administration. The animals were separated into five groups with five mice per cage (n = 5 mice). The first group of animals was injected with a single dose of MPTP dissolved in saline solution at a concentration of 20 mg/kg. The second group was injected with the same amount of saline solution, the third group with the Cdc42 inhibitor (ML141) dissolved in corn oil at a concentration of 10 mg/kg, the fourth group with the same amount of corn oil, and the final group was injected with ML141, 30 minutes prior to MPTP injection. No anesthesia or analgesia method was indicated or recommended. After 72 hours, animals were euthanized, intracardially perfused with Tyrode’s solution and fixed with 4% PFA. Brains were extracted and post-fixed in 4% PFA during 48h. After post fixation, brains were stored in PBS and NaN_3_, until further sectioning.

#### MPTP intoxication and passive immunotherapy with CD16/32 neutralizing antibodies

To see the effect of blocking CD16/32 Fcγ receptors in a parkinsonian model, three-week-old C57BL/6 male mice (n = 25) were commercially obtained (Janvier Labs). As described in the previous section, in experiments using MPTP, the animals were kept and maintained in special and isolated conditions due to the toxicity of MPTP. Six cages of five animals each were used (n = 5 mice). The first group was injected with MPTP dissolved in saline solution, the second group with the same volumetric amount of saline solution. A third group was injected with CD16/32 neutralizing monoclonal antibodies (Ultra-LEAF Purified Anti-mouse CD16/32 Antibody, Biolegend, San Diego, CA, USA) in a volume of 200 µL/mouse (10 mL/kg) 24 hours prior to MPTP administration (20 mg/kg). A fourth group was injected with a control isotype antibody (Ultra-LEAF Purified Rat IgG2a, κ Isotype Ctrl Antibody), also in a volume of 200 µL/mouse (10 mL/kg) 24 hours prior to the MPTP injection. The fifth group, as control of the antibody, was injected with a single dose of CD16/32 blocking antibody 24 hours before the injection of saline solution (n=3 mice), and the final group, labeled as control Isotype, an injection of the isotype rat antibody was injected 24 hours before an injection of saline solution (n=2 mice). After 72 hours, animals were euthanized by intracardiac perfusion of Tyrode’s solution and fixed with 4% PFA. Brains were then extracted and post-fixed during 48h in 4% PFA. After post fixation, brains were stored until further sectioning in tubes with PBS and NaN_3_.

#### Mouse brain sectioning

Stored brains were prepared for sectioning in a conventional mouse brain matrix to obtain two blocks. One block containing the striatum (4 mm from Bregma,) and another block containing the mesencephalon (-4 mm from Bregma) according to the mouse brain in stereotaxic coordinates atlas^42^. After removing the olfactory bulbs, each brain block was sectioned by vibratome (Leica Microsystems, Wetzlar, Germany). Six series of 40-µm sections were obtained, setting the vibratome at a speed of 0.225 mm/s and a vibrating frequency of 60 Hz. Once sectioned, serial sections of the brain tissue were stored in PBS with NaN_3_ until DAB immunohistochemistry or multicolor immunofluorescence was performed.

### Cell cultures

#### Cell lines

To study the behavior of microglial and dopaminergic cells in a PD-like neuroinflammatory scenario, we used cell lines BV2 microglia and PC12 dopaminergic cells. BV2 cells were maintained at 37°C and 5% CO_2_ in Roswell Park Memorial Institute (RPMI) medium (Sigma Aldrich; Saint Louis, MO, USA), supplemented with 10% heat-inactivated fetal bovine serum (FBS) and 0.1% penicillin-streptomycin (P/S) whereas PC12 cells were maintained at 37°C and 5% CO_2_ in Dulbeccos Modified Eagle Medium (DMEM) (Sigma Aldrich; Saint Louis, MO, USA), supplemented with 7% heat-inactivated fetal bovine serum (FBS), 7% horse serum (HS), 1.14% HEPES 1M pH=6.8 and 0.2% P/S.

#### Treatment with interferon-gamma and lipopolysaccharide

To see the effect of proinflammatory inductors, microglial cells were seeded on coverslips in 24-well culture plates at a density of 20,000 cells per milliliter (cells/mL). Cells were then exposed to increasing concentrations of interferon-gamma (IFN-γ) (0.05, 0.5, 5, 25, 50, 75 and 100 ng/mL) or combined with proinflammatory stimulus lipopolysaccharide (LPS) (Sigma-Aldrich, St. Louis, MO, USA) at a concentration of 100 ng/mL. After 24 hours (day *in vitro* 1) the supernatant was collected for further studies. New fresh medium was added and 24 hours later (day *in vitro* 2) supernatant was collected then the cells were fixed with 4% paraformaldehyde (PFA) for immunostaining.

#### Microglial priming

Regarding understanding microglial priming, the order of the proinflammatory stimulus was alternated to see if there were variations in the level of activation of BV2 cells. Microglial cells were seeded on coverslips in 24-well culture plates at a density of 20,000 cells/mL, 24 hours later adding either IFN-γ, LPS or the combination of both to the medium (IFN-γ at 0.5 ng/mL and LPS at 100 ng/mL). After 24 hours, the supernatant was collected for further studies and new fresh medium with LPS, IFN-γ or the combination of both, was added to each well. After 24 hours, supernatant was collected, and the cells were fixed with 4% PFA for immunolabeling.

#### Detection of microglial activation by Griess assay

The colorimetric analysis Griess assay (Sigma-Aldrich, St. Louis, MO, USA) was used as a control to detect the presence of nitrite compounds in the supernatant, in this case, product of the activation of BV2 cells. A calibration curve was determined using NaNO_2_ (sodium nitrite) in a range between 0.78 µM and 100 µM. Griess reactive (100 µL/well) was added to a 96-microwell plate with 100 µL of each of the samples. After 15 minutes of incubation at room temperature, absorbance was read at 540 nm and the values were interpolated into the calibration curve to obtain the concentration of nitrites in the supernatants.

#### Analysis of BV2 and PC12 interactions

To see the cell-cell interaction and the effect of the activation between microglial and dopaminergic cells, a co-culture with BV2 and PC12 cells was set up. BV2 cells were seeded in a 24-well plate at a concentration of 300,000 cells/mL. Afterwards, the cells were stimulated with the combination of LPS 100 ng/mL plus IFN-γ 0.5 ng/mL and 24 hours later the medium with the treatment was removed from the wells. Previously trypsinized PC-12 cells were seeded on top of the BV2 cells with fresh DMEM medium at a density of 1,200,000 cells/mL. The co-culture was then fixed with 4% PFA after 10, 20 and 30 minutes and 1, 4 and 10 hours of interaction. Fixed cells were preserved in PBS to later perform multicolor immunocytofluorescence. Alternatively, because prior priming with IFN-γ, followed by LPS, reaches a higher level of activation, we defined a sequential procedure. After 24 hours of incubation BV2 cells were first stimulated with IFN-γ at a concentration of 0.5 ng/mL. Twenty-four hours later BV2 cells were challenged with LPS at 100 ng/mL. After 24 hours of incubation, PC-12 cells were seeded on top of BV2 cells at a density of 200,000 cells/mL. The co-culture was fixed with 4% PFA at 10, 20 and 30 minutes and 1, 4, 10 and 24 hours of interaction. Fixed cells were preserved in PBS until performing immunocytofluorescence with multiple markers.

#### Cdc42 inhibition in BV2 and PC12 cell co-cultures

To analyze how the inhibition of the protein Cdc42 affects the preservation of dopaminergic cells, the inhibitor ML141 (Cdc42/Rac1 GTPase Inhibitor, Calbiochem, San Diego, CA, USA) was used in BV2/PC12 co-cultures. This experiment was divided in two scenarios. In the first, inhibition with ML141 was done before activation with IFN-γ and LPS and in the second, inhibition was done after activation. According to the aforementioned protocol, BV2 cells were seeded at a concentration of 50,000 cells/mL and incubated at 37°C and 5% CO_2_. For inhibition before activation, ML141 was added at a concentration of 10 µM, diluted in DMSO, with new fresh medium to the BV2 cells after 24 hours of being seeded. One hour later, the medium was removed and new RMPI medium was added with IFN-γ at 0.5 ng/mL. After 24 hours, the medium was removed and LPS 100 ng/mL was added, according to the priming protocol described above. PC12 cells were seeded on top of the BV2 cells at a concentration of 200,000 cells/mL and after one hour of incubation cells were fixed with 4% PFA and preserved in PBS to conduct immunocytofluorescence.

#### Inhibition of CD16/32 in BV2 and PC12 cells interactions

As described for Cdc42, we followed the same inhibition procedure in pre-activation and post-activation scenarios. Cells were treated with CD16/CD32 blocking antibody (Ultra-LEAF Purified Anti-mouse CD16/32 Antibody, Biolegend, San Diego, CA, USA), to prevent phagocytosis, or its isotype (Ultra-LEAF Purified Rat IgG2a, κ Isotype Ctrl Antibody). BV2 and PC12 cells were seeded proceeding as described above and the cells were activated with IFN-γ and LPS before or after being treated with either CD16/32 neutralizing antibody or the isotype at a concentration of 10 µg per 10^6^ cells. After one hour of interaction, cells were fixed and preserved in PBS.

### Brain tissue processing and immunolabeling

#### Immunohistochemistry for human samples

Thionine staining was performed to analyze the anatomical structure of the hemi-mesencephala, detect the nuclei of the cells, and visualize the cytoplasm of neuromelanin-containing neurons of the SNpc. Immunohistochemistry by diaminobenzidine (DAB) detection was performed on serial brain sections to visualize markers of neuroinflammation and phagocytic capacity of microglial cells, including FcγR CD16 and CD32 (**Table 2**). After PBS washing, a solution of H_2_O_2_ 0.3% was added to the tissue sections for 15 minutes to inactivate the endogenous peroxidase. Then nonspecific binding sites were blocked by incubating the sections with 0.5% Triton X-100 with 10% horse serum (HS) (Sigma-Aldrich; St. Louis, MO, USA) for one hour. After blocking, primary antibodies were used during 48 hours, diluting the antibodies in Trizma base saline (TBS)-0.5% Triton X-100, 1% HS and 0.1% NaN_3_ (Trizma Base Sigma-Aldrich; St. Louis, MO, USA). Then secondary detection was carried out with a specific biotinylated antibody (Table 2) diluted in 0.5% Triton X-100 with 1% HS without NaN_3_. Biotin of the secondary antibody was detected using the Vectastain Elite ABC horseradish peroxidase method (Vector Laboratories; CA, USA) following the manufacturer’s instructions. After DAB detection, sections were placed on gelatinized glass slides and were dehydrated through graded ethanol solutions (70, 80, 90 and 100%) and submerged in xylene before placing the coverslips with appropriate mounting media (Xylene substitute mounting medium. Thermo Fisher Scientific; Waltham, MA, USA).

**Table 2.** List of primary and secondary antibodies used in immunofluorescence and immunohistochemistry.

| Primary antibodies |  |  |  |
| --- | --- | --- | --- |
| Antibody | Source | Dilution | Company [Reference] |
| anti-TH | sheep polyclonal | 1:500 | Sigma [ab152] |
| anti-Iba1 | rabbit polyclonal | 1:500 | Wako [019-19741] |
| anti-Cd11b | rat monoclonal | 1:100 | Bio-rad [MCA711G] |
| anti-CD16/32 | rat | 1:250/200 | BD Pharmigen [553142] |
| anti-CD16 | goat | 1:50 | RD Systems [AF1597] |
| anti-CD32 | mouse | 1:100 | Bio-rad [MCA1075] |
| anti-CD32 | mouse | 1:100 | Biosource [MBS249188] |
| anti-Cdc42 | mouse monoclonal | 1:500 | Cytoskeleton [ACD03-A] |
| anti-F4/80 | rat | 1:50 | Bio-rad [MCA497] |
| anti-MAP-2 | chicken polyclonal | 1:500 | Abcam [ab5392] |
| anti-NeuN | mouse monoclonal | 1:500 | Merck [MAB377] |
| Secondary antibodies |  |  |  |
| Antibody |  | Dilution | Company |
| Biotinylated donkey anti sheep |  | 1:1000 | Jackson [713-065-003] |
| Biotinylated donkey anti goat |  | 1:200 | Jackson [705-065-147] |
| Biotinylated goat anti rabbit |  | 1:1000 | Dako [BA-1000] |
| Alexa Fluor 555 goat-anti rabbit |  | 1:1000 | ThermoFisher [A21429] |
| Alexa Fluor 488 donkey-anti sheep |  | 1:1000 | ThermoFisher [A11015] |
| Alexa Fluor 488 goat anti-rat |  | 1:1000 | ThermoFisher [A11006] |
| Alexa Fluor 555 goat anti-rat |  | 1:1000 | ThermoFisher [A21434] |
| Alexa Fluor 488 goat anti-mouse |  | 1:1000 | ThermoFisher [A21121] |
| Alexa Fluor 488 goat anti chicken |  | 1:1000 | ThermoFisher [A11039] |

#### Immunohistochemistry for mouse brain samples

Immunohistochemistry by diaminobenzidine (DAB) detection was also performed on mouse brain sections to visualize microglial cells and dopaminergic neurons using anti-Iba-1 and anti-TH antibodies respectively (**Table 2**). Antigen retrieval was conducted with a pre-treatment with citrate (10mM, pH 6, 60°C) for 20 minutes. The same protocol described above for human samples was done for mouse brain sections. Variations to this procedure were made for immunodetection of CD16/32 positive cells in the tissue, including the primary antibody concentration, and a higher citrate temperature (10mM, pH 6, 80 °C) (**Table 2**).

#### Immunohistofluorescence for mouse brain samples

To detect changes in microglia due to neuroinflammation and its effect on dopaminergic neurons, anti-Iba-1 antibody was used to detect and visualize microglial cells in combination with anti-TH antibodies to detect dopaminergic neurons. Anti-CD16/32 was used to mark phagocytic microglia/macrophages. Antigen retrieval was done routinely as described before with citrate at 60 °C and blocking of nonspecific antibody binding sites during one hour with 10% HS. Primary antibodies were diluted in TBS-0.5% Triton X-100, 1% HS and 0.1% NaN_3_ and incubated for 48 hours. Suitable secondary antibodies (Table 2), diluted in TBS-0.5% Triton X-100, 1% HS and 0.1% NaN_3_, were used accordingly and incubated overnight. Finally, DAPI (1:1000; Life technologies; Carlsbad, CA, USA) was used to stain the nuclei of the cells. Sections were mounted on glass slides and coverslipped using an appropriate antifading reagent (Prolong Gold, Invitrogen; Carlsbad, CA, USA).

### Cell culture processing and immunolabeling

#### Immunocytofluorescence for cell lines

In the case of the IFN-γ/LPS treatment, microglial cells were stained to visualize marker expression and its morphological variations. Cells were then treated for permeabilization and antigen retrieval with PBS with saponin 0.02% for seven minutes. After washing, a blocking solution was added (PBS with 0.01% saponin, 10mM glycine) for 15-minute incubation at room temperature. A more concentrated blocking solution (PBS with 0.01% saponin, 10 mM glycine, 5% BSA) was then added for one hour at room temperature. Primary antibodies were diluted in PBS with 0.01% saponin, 1% BSA (Sigma-Aldrich, St. Louis, MO, USA), and placed in a humidity chamber overnight at room temperature. After washing with PBS, appropriate secondary antibodies (Table 2) were added for 45 minutes. Phalloidin staining was also performed to visualized F-actin cytoskeleton (AlexaFluor 555 Phalloidin ThermoFisher). Nuclei were counterstained with DAPI (1:1000) for five minutes before washing with PBS. Finally, antifading reagent was used to mount the coverslips on glass slides (Prolong Gold, Invitrogen; Carlsbad, CA, USA).

### Microscopy analysis, stereology, and quantifications

#### Microscopy, stereological quantification in human brain samples

Sections were examined and analyzed with a conventional light microscope (Eclipse 80i microscope, Nikon, Tokyo, Japan), taking ten random fields of dorsal and ventrolateral areas of the SNpc of each representative section with a 20X objective and a digital camera connected to the microscope (DXM 1200F Digital Camera, Nikon; Tokyo, Japan). Sampling fields were processed for stereological quantification to measure cell density applying appropriate software tools (Image J version 1.47, NIH, USA).

#### Microscopy, stereological quantification in mouse brain samples

A bright field microscope (Eclipse 80i microscope, Nikon; Tokyo, Japan) with a connected digital camera (DXM 1200F Digital Camera, Nikon, Tokyo, Japan) was also used for taking the required sampling. Based on the anatomy of the mouse brain defined in the Mouse Brain Atlas^42^, motor cortex and striatum were analyzed. Six randomized dissector photo-samples of the motor cortex and six of the striatum were taken with 20X objective. The area occupied by Iba-1 was measured with the required software tools (Image J version 1.47, NIH, USA) establishing a threshold for every stained cell of the field.

#### Microscopy, stereological quantification in cell lines

The number of cells expressing Iba-1 or TH was also quantified by stereological methods in the experiments of increasing IFN-γ concentration and co-cultures. Each coverslip was systematically sampled using a fluorescence microscope (Nikon Eclipse 90i, Nikon, Tokyo, Japan) attached to a digital camera (DXM 1200F) and software (ACT-1 version 2.70, Nikon corporation) with a 20X objective, taking 10 sample pictures per coverslip, per condition. Once the pictures were taken, Iba-1 and TH positive cells were quantified with cell counter image-analysis software (Image J version 1.47, NIH, USA), as well as their interactions and phagocytic events, always following stereological criteria: each frame was considered as a physical dissector and only particles in focus in one Z optical plane were quantified. The area occupied by Iba-1 and DAPI was also measured with the aforementioned software (Image J) establishing a threshold for every fluorophore. The threshold was critical to include all the positive material and exclude the background.

### Confocal imaging and high-resolution analysis

#### Confocal microscopy, stereological analysis in mouse brain sections

To properly visualize detailed changes in the quantity of dopaminergic neurons, sections of the SNpc, identified by the Mouse Brain Atlas criterion^42^ were imaged with high resolution in a confocal laser-scanning microscope (LSM 700; Carl Zeiss, Oberkochen, Germany), using a Plan-Apochromat 20x/0.8 Ph2 M27 objective lens (Carl Zeiss; Oberkochen, Germany), and processed with the proper capture software (ZEN 2010 Carl Zeiss; Oberkochen, Germany). At the 20X objective level, the optical dissector was able to cover the extent of the SNpc, therefore, two to four serial sections were imaged for both hemispheres and the number of dopaminergic neurons was quantified (with Image J software), obtaining the average number of cells per SNpc, per animal. A magnification of 1X was applied to the scanning field using the 20X objective to achieve higher definition of microglial cells. The image analysis software and plugin (Image J software) allowed the transformation of images to black and white, and with the adequate threshold, the area of every cell was measured and calculated. In addition, representative images were taken with a confocal laser-scanning microscope (LSM 700; Carl Zeiss, Oberkochen, Germany) and a Plan-Apochromat 40x/1,3 Oil DIC M27 objective lens (Carl Zeiss; Oberkochen, Germany), and processed with specialized software (ZEN 2010 Carl Zeiss; Oberkochen, Germany). A series range for each section was determined by setting an upper and lower threshold using the Z/Y position for the Spatial Image Series setting covering the cell height. Then, the microscope captured pictures at a fixed distance over the entire volume of the cells. Stacks of the images were taken with an optical section interval, from 0.5 μm. Pictures were analyzed with appropriate image analysis software (Imaris, Bitplane) to visualize three-dimensional structures at high resolution. Representative microglia were processed for isosurface generation to visualize the cell size and arborization. Phagocytic events were processed for three-dimensional visualizations with isosurfaces to visualize the phagosomes. And lastly, confocal images were processed for relative fluorescence analysis by displaying the plot profile at the region of interest (Image J).

#### Analysis of segregation of CD16/32 at the phagocytic synapse in vitro

High-resolution and three-dimensional representative images of microglia-dopaminergic cell interactions from the co-culture were taken with a confocal microscope (ZeissExaminer D1 AX10). Using a digital cropping zoom, captured with confocal analysis software (ZEN2010), a high-definition series range for each interaction was set by determining an upper and lower threshold using the Z/Y position for spatial image series with the 40x immersion oil objective. Image stacks were taken with a 0.5-μm optical section interval. Sets of images were analyzed using image analysis software (Image J, version 1.52v, NIH, USA). To study the localization of the phagocytic receptor CD16/32 in the interaction between BV2/PC12 cells, we set a line covering the interacting interface of the BV2 cell in contact with the PC12 cell. We measured the relative fluorescence of CD16/32 and the TH of the interacting target cell along the adapted line of interface, obtaining a plot profile. Early-stage phagocytosis events and phagocytic synapses were captured and analyzed in detail; to analyze the CD16/32 fluorescence segregation in the phagocytic cup, we analyzed representative interaction events. We assigned the highest number of the relative fluorescence from each image a value of 100% and calculated the percentages in the different areas of the interface in relation to that value. We segmented the established line of interaction in two parts considering the position of PC12 and the establishment of phagocytic interactions. “Periphery” was considered at the limits of the intercellular interactions of BV2 microglia with PC12 DA cells, and “center” where the PC12 cells were indeed interacting with BV2. The average of calculated percentages for each segment of the profiled line were used to analyze the segregation of CD16/32.

### Statistics

All the data were expressed as the mean with the SEM (standard error of the mean). Regarding the cell cultures and the Griess assay, every experiment was performed at least five times. Normal distribution of the data was tested with the Shapiro-Wilk and Pearson normality test, the P value being greater than the α value (α=0.05) to prevent discarding the gaussian distribution of the data. Once normality was tested, a one-way ANOVA and multiple comparisons were conducted to compare the mean of every treatment, both among treatments and, especially, against the control. Tukey’s post hoc test was also performed, with α=0.05 and P value smaller than α to accept a significant difference. For in vivo experiments, for the intraperitoneal administration of ML141, every group counted with n=5. One of the groups did not follow a normal distribution; therefore, logarithmic modification was applied to every data set to analyze the data with one-way ANOVA with Tukey’s multiple comparison post hoc. Concerning the MPTP experiments, the quantification data, both the number of neurons and the area of Iba-1 were tested with Pearson’s, Shapiro-Wilk and Kolmogorov-Smirnov, indicating that normality should not be dismissed. Therefore, assuming variances were equal, comparison was done by one-way ANOVA and the LSD (least significant difference) post hoc. In all cases, P ≤ 0.05 was considered significant. All statistical analyses were performed using Prism 6 (GraphPad Software; La Jolla, CA, USA) and SPSS (IBM SPSS Statistics for Windows, Version 21.0. Armonk, NY: IBM Corp) software.

## Acknowledgements

This work was supported by grants from the Spanish Ministry of Sciences and Universities and the European Regional Development Fund (*Fondo Europeo de Desarrollo Regional,* FEDER) (Reference grant (PGC2018-096003-B-I00), the Spanish Ministry of Science and Innovation (Reference grants PID2021-128717OB-I00, and PRE2022-101261, Fellowship for IFB), the Generalitat de Catalunya (Reference grant, 2014 SGR-984, Fellowship for PCR), the Parkinson’s and Movement Disorders Foundation, USA (PMDF’2016 Grant Program), and the American Parkinson’s Foundation-APDA Summer Student Fellowship 2019 to EC and CGM. We would also like to thank all the personnel from the Administration and Technical Laboratories of the *Institut de Neurociències*, for the help provided at the *Universitat Autònoma de Barcelona*, very especially to the outstanding technicians at the Cell Culture Unit, Cristina Gutierrez, Antonio Cambero and Neus Ontiveros, the Histology Lab, Mar Castillo and Anna Torrent, and the Microscopy Core, Meritxell Roig-Martínez (coathor of this work), Núria Barba and Saioa Mendizuri.

## List of grants and funding agencies

Ministerio de Ciencia Innovación y Universidades (PGC2018-096003-B-I00)

Ministerio de Ciencia e Innovación (PID2021-128717OB-I00)

Ministerio de Ciencia e Innovación (PRE2022-101261) Fellowship for IFB

Generalitat de Catalunya, 2014, SGR-984, Fellowship to PCR

The Parkinson’s and Movement Disorders Foundation, USA. PMDF’2016 Grant Program

American Parkinson’s Foundation-APDA Summer Student Fellowship 2019 to CGM

American Parkinson’s Foundation-APDA Summer Student Fellowship 2019 to EC

