## Supplementary figures and images for "Microglial low-affinity FcγR mediates the phagocytic elimination of dopaminergic neurons in Parkinson’s disease degeneration"

### Supplementary figure 1

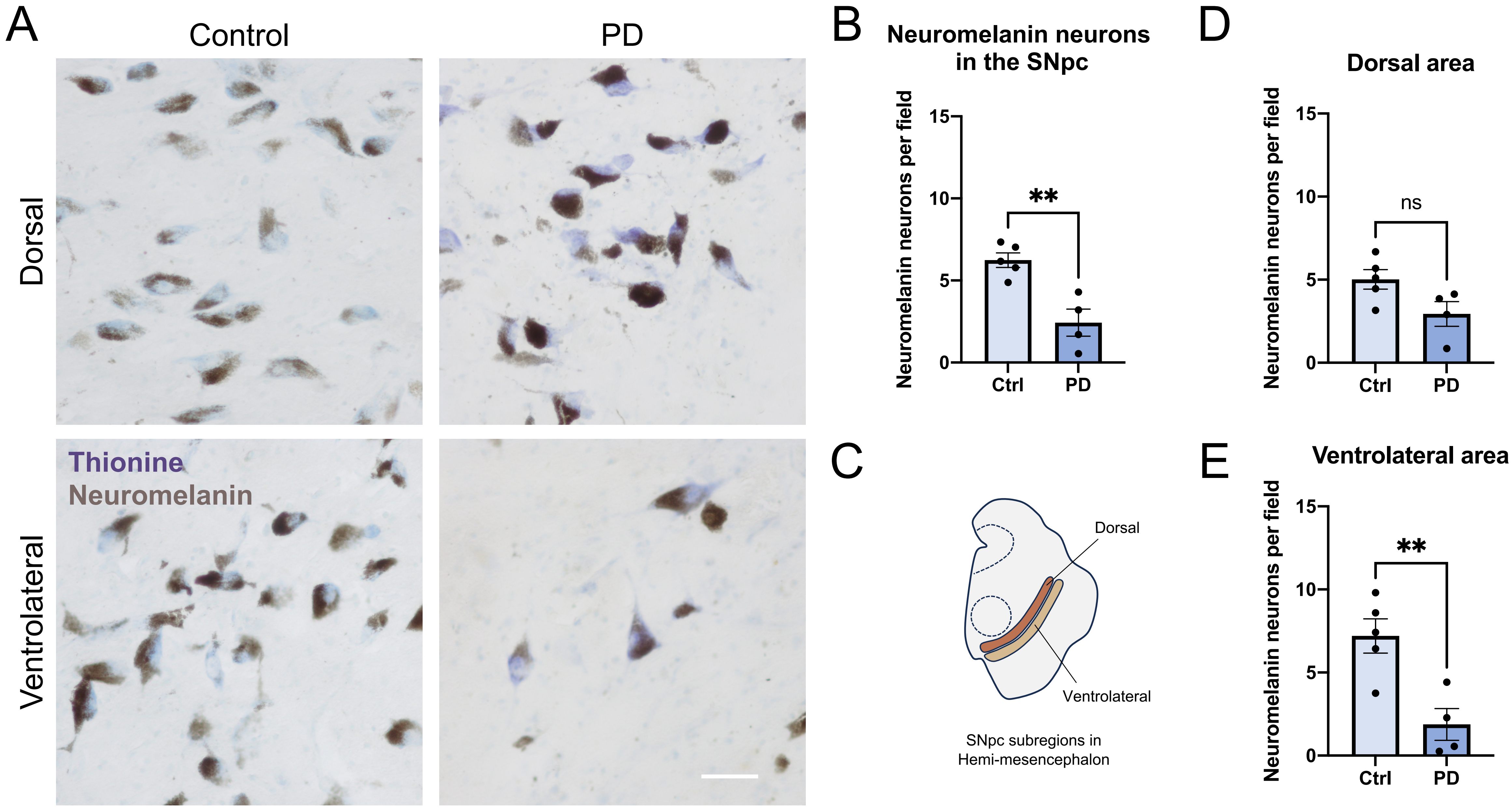


**Supplementary figure 1**

### Supplementary figure 2

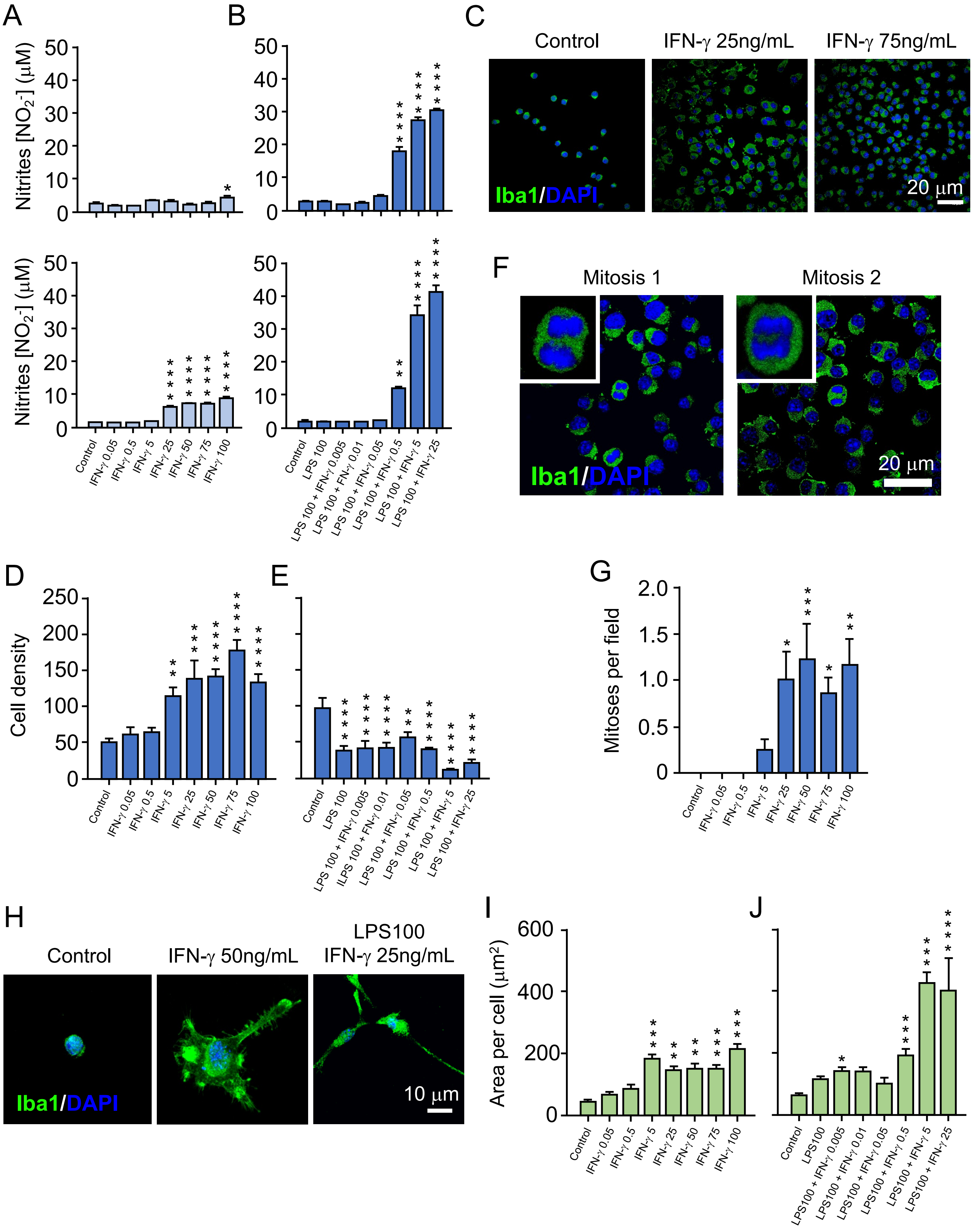


**Supplementary figure 2**

### Supplementary figure 3

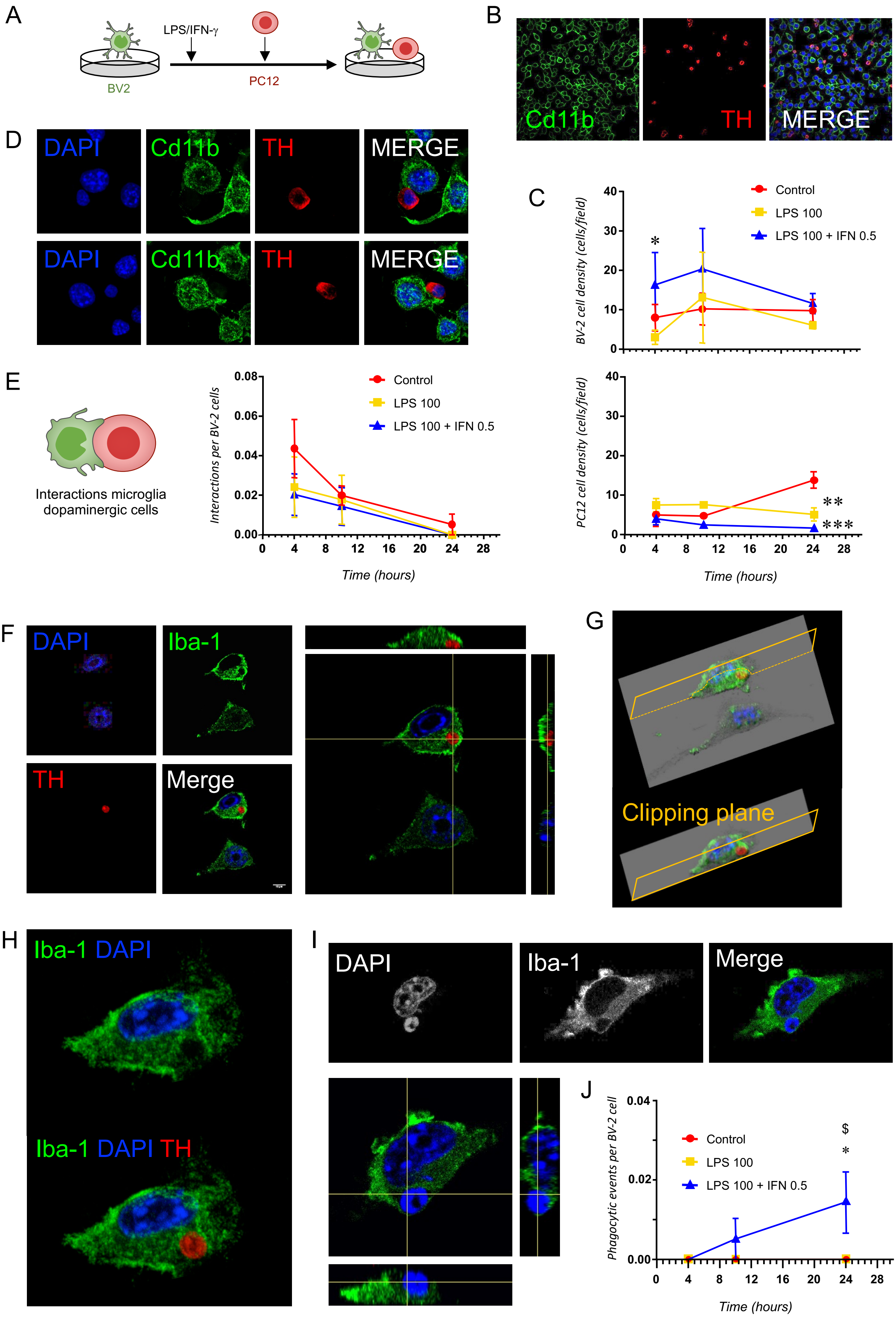


**Supplementary figure 3**

### Supplementary figure 4

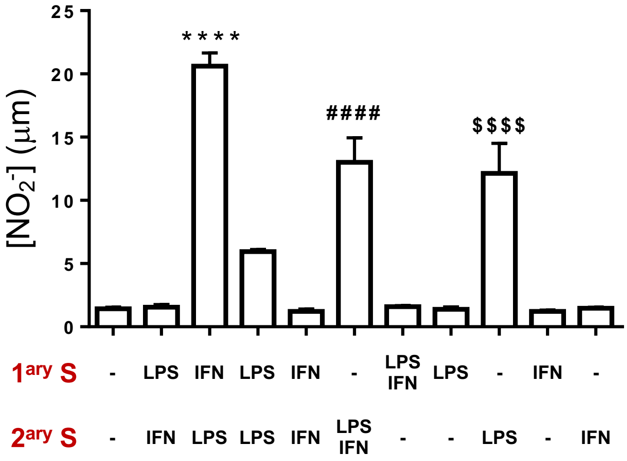


**Supplementary figure 4**

### Supplementary figure 5

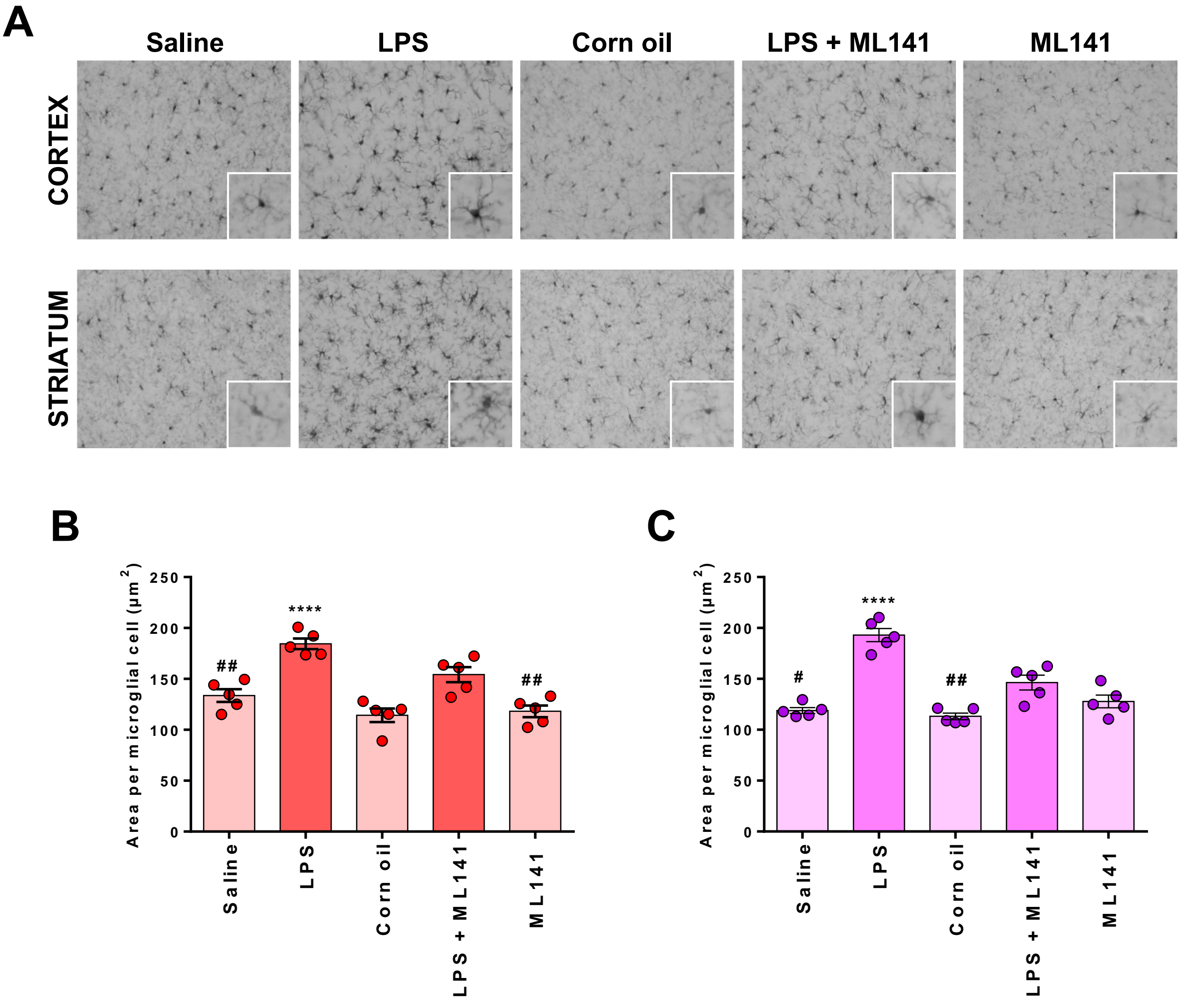


**Supplementary figure 5**
